# Microbial reaction rate estimation using proteins and proteomes

**DOI:** 10.1101/2024.08.13.607198

**Authors:** J. Scott P. McCain, Gregory L. Britten, Sean R. Hackett, Michael J. Follows, Gene-Wei Li

## Abstract

Microbes transform their environments using diverse enzymatic reactions. However, it remains challenging to measure microbial reaction rates in natural environments. Despite advances in global quantification of enzyme abundances, the individual relationships between enzyme abundances and their reaction rates have not been systematically examined. Using matched proteomic and reaction rate data from microbial cultures, we show that enzyme abundance is often insufficient to predict its corresponding reaction rate. However, we discovered that global proteomic measurements can be used to make accurate rate predictions of individual reaction rates (median *R^2^* = 0.78). Accurate rate predictions required only a small number of proteins and they did not need explicit prior mechanistic knowledge or environmental context. These results indicate that proteomes are encoders of cellular reaction rates, potentially enabling proteomic measurements *in situ* to estimate the rates of microbially mediated reactions in natural systems.

**Significance:** One of the most basic phenotypes of a microbe is its set of associated reaction rates, but quantifying these rates *in situ* remains extremely challenging, especially in natural systems. We used molecular data and statistical models to estimate microbial rates in steady state cultures. We found that many reaction rates are highly predictable using proteomic data, though single proteins are typically not informative for their associated reaction rates. This result suggests that gene expression data from complex microbial communities could be used to estimate *in situ* reaction rates, providing new clues into the lives and environmental function of microbes.

## Introduction

Microbes dramatically shape their environments via the multitude of reactions they perform. For example, a subset of microbes fix dinitrogen gas into ammonia, supplying bio-available nitrogen and maintaining the productivity of the global biosphere (1). In the human gut, bacteria produce and consume hydrogen sulphide, processes that are implicated in several gut-related diseases (e.g., 2). In such examples, a fundamental challenge is to quantify these rates. Successfully quantifying individual reaction rates *in situ* is a first step towards determining the environmental conditions that lead to higher or lower rates and, ultimately, the mechanistic underpinnings of the reactions. With taxonomic resolution, such data would provide significant new insight into the ecological dynamics of microbially mediated systems, including the global carbon cycle and the human microbiome.

Although measuring *in situ* reaction rates can be extremely challenging, measuring the enzymes that mediate such rates is much easier. For example, directly inferring bacterial metabolic processes in the deep sea, or in the human body, is significantly more challenging than collecting samples and characterizing the metatranscriptome or metaproteome. Advances in sequencing technologies and mass spectrometry have contributed to this accessibility and have yielded an unprecedented window into *in situ* microbial activities. If data on enzyme abundances could be used to quantitatively estimate rates, there are two main advantages: 1. Because transcripts and proteins contain taxonomic information, community-level biogeochemical rates could be deconstructed into taxon-level contributions. 2. The high-throughput methods used to evaluate enzyme stocks could be leveraged to dramatically increase the scale of rate estimates, especially in natural environments.

Several approaches have combined gene expression data with mechanistic models to make predictions about reaction rates (3–9). However, these approaches require *a priori* knowledge of the biochemical network, and reactions with unclear enzymatic underpinnings are sometimes excluded. For example, extracellular superoxide production is an important biogeochemical process, but the mechanistic bases are not well-understood in many taxa (10, 11). Other studies focused on natural environments have incorporated other data sources, e.g., elemental stoichiometry, but they similarly require prior biochemical knowledge (12, 13). Overall, it is unclear if these types of mechanistic models can be readily extended to all reactions of interest.

Reaction rate information may also be encoded in transcriptomes and proteomes (without the requirement for constraints from genome-scale metabolic models). For example, Kochanowski *et al* demonstrate that the catabolic repressor activator Cra in *Escherichia coli* can be used as a “flux sensor” because its transcription factor activity is negatively correlated with glycolytic flux (14). (Note that we use the word “rate” instead of “flux” throughout, because flux is typically normalised by area in environmental science.) Pioneering work suggests that it is possible to infer specific biogeochemical rates using even single proteins, particularly when substrate concentrations are constrained or assumed (15, 16). However, we do not know if this assumption can be generalised to other reaction rates, particularly when we do not have prior information about relevant substrate concentrations. Indeed, a simple toy model of metabolic pathways suggests that enzyme abundance does not always covary with their cognate rates (Box 1).

In this contribution, we systematically examined empirical relationships between enzyme abundances and intracellular reaction rates (hereafter termed simply as “rates”). Using a range of datasets from microbial cultures in steady state conditions, we estimated covariation of enzyme abundance and rates. We found that many protein abundances are only weakly correlated with the rates of their associated reactions, indicating that single proteins alone are unlikely to be informative for inferring reaction rates in natural microbial communities. Our finding challenges a common tacit assumption in experimental and observational work that a change in protein abundance corresponds to a change in the process this protein mediates. However, we discovered that many rates are highly predictable when using additional proteomic context, such as other proteins within the given pathway, or the entire proteome. We conclude that proteomic and transcriptomic data present a promising avenue for taxon-specific quantification of *in situ* reaction rates in natural microbial systems.

### Box 1

#### Pathway structures complicate enzyme rate relationships

If many reaction rates were simply controlled by modulating the abundance of the associated enzyme, then knowledge of a single protein’s concentration might be sufficient to predict a cognate rate. A change in the abundance of a transcript or protein could, at the very least, be used to interpret a change in the corresponding process that such protein mediates. A naive expectation is from first order enzyme kinetics, *v* = *k* ⋅ *e* ⋅ *s*, where *v* is the rate of a reaction*, e* is the enzyme concentration, *s* is the substrate concentration and *k* is the first-order rate constant (Fig. 1a). If *e_1_* increases, there is an increase in the associated reaction rate (*v_1_*) assuming that the enzyme concentration is *independently* modulated (i.e., without any simultaneous changes to the kinetics or substrate concentration). As an example of how this assumption can be easily violated, consider a branched pathway where two enzymes compete for a single shared substrate. The substrate concentration is now a dynamic variable, as opposed to an imposed parameter as above (Fig. 1b). If concentrations of *e_1_* and *e_2_* are correlated (e.g., *e_2_* is a function of *e_1_*, Fig. 1b) and the substrate input rate (*C*) is fixed, then an increase in *e_1_* could correspond with a decrease in its cognate reaction rate, *v_1_* (Fig. 1b). Real metabolic networks are much more complicated than this simple branched pathway, so relationships between rates and enzyme concentrations are non-trivial. Another route for breaking this *independent modulation* assumption is via changes to the first-order rate constant, *k*, for example via post-translational modifications (e.g., 17).

**Figure 1.**
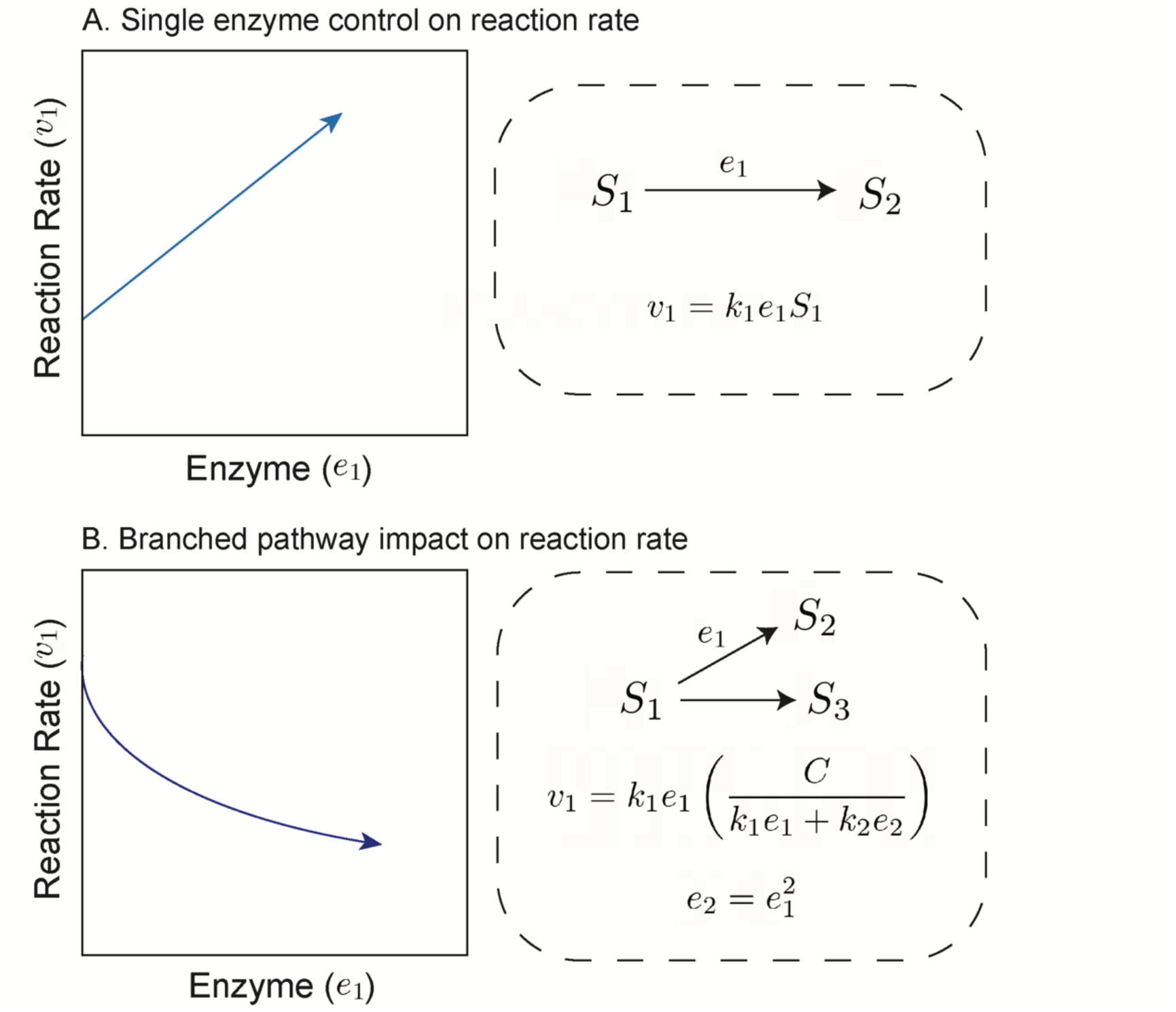
Covariation between enzyme abundance and reaction rate. (A) Single-enzyme control on reaction rate suggests a simple relation between enzyme concentration and its associated reaction rate, where *S_1_* is a substrate, *S_2_* is a product, *k_1_* is the first-order rate constant, and *e_1_* is the enzyme concentration. Here, *S_1_* and *k_1_* are assumed to be constant. (B) A branched pathway changes enzyme-reaction rate relationships by indirectly modulating *S_1_*: here we set *e_1_* as a function of *e_2_*, which are enzymes that compete for the same substrate *S_1_*. *C* is the rate of supply of *S_1_*. Assuming that *C* is constant and *S_1_* is not, increasing *e_1_* leads to a decrease in *v_1_*.

The branched pathway model indicates that information about the proteomic context (e.g., enzyme *e_2_* in Fig. 1b) or about substrate input rate (reflected in the constant *C*, Fig. 1b), could be leveraged to constrain reaction rate estimates. Variance in a given reaction can arise from at least three sources representing different scales of metabolic control: 1) the focal enzyme concentration (that mediates a given reaction rate, e.g., *e_1_* in Fig. 1b), 2) within-pathway proteins (i.e., the local neighbourhood of proteins, e.g., both *e_1_* and *e_2_* in Fig. 1b), and 3) the global context of proteins; those beyond the local pathway that are not explicitly depicted in Fig. 1. These three scales provide an outline for our analyses in this work. We begin by systematically examining single-protein to rate covariation, then examine how single rates can be predicted using within-pathway proteins, and finally use proteomes to predict individual rates.

## Results

### Single proteins have diverse statistical relationships with their associated rates

We investigated empirical relationships between intracellular rates and their associated protein abundances. To systematically examine protein-to-rate relationships, we first utilized a dataset of *Saccharomyces cerevisiae* in steady-state growth under diverse conditions (15 conditions, ref. 18). Specifically, the yeast cultures were grown in chemostat with 5 different dilution rates of three different media types (phosphate-limited, ammonia-limited, and glucose-limited).

Protein abundance was quantified using isotope-labelled mass spectrometry and is normalised to total cellular proteins. Reaction rates (in units of mmol ⋅ hour^-1^ ⋅ mL cell volume^-1^) were derived by metabolomic measurements coupled with a cellular scale metabolic model (18, 19). Importantly, no proteomic or transcriptomic data were used to derive reaction rates. Both proteomic measurements and derived rates were previously published. For this initial analysis, we only considered reactions with a unique dedicated protein, which resulted in 46 protein-to-rate relationships.

In general, we found a diverse set of statistical relationships between individual protein abundances and their associated rates. Our systematic assessment suggests that the naive expectation of covarying rate and protein abundance is not commonly observed. For example, saccharopine dehydrogenase (LYS9), an enzyme involved in lysine biosynthesis, shows a simple positive relationship between the log-transformed protein abundances and reaction rates (Fig. 2A), with a high Spearman correlation coefficient (0.76). However, other proteins exhibited unexpected relationships with their associated rates, such as monomeric glyoxylase I (GLO1), which displayed a negative relationship (Spearman correlation coefficient = −0.81; Fig. 2B).

**Figure 2.**
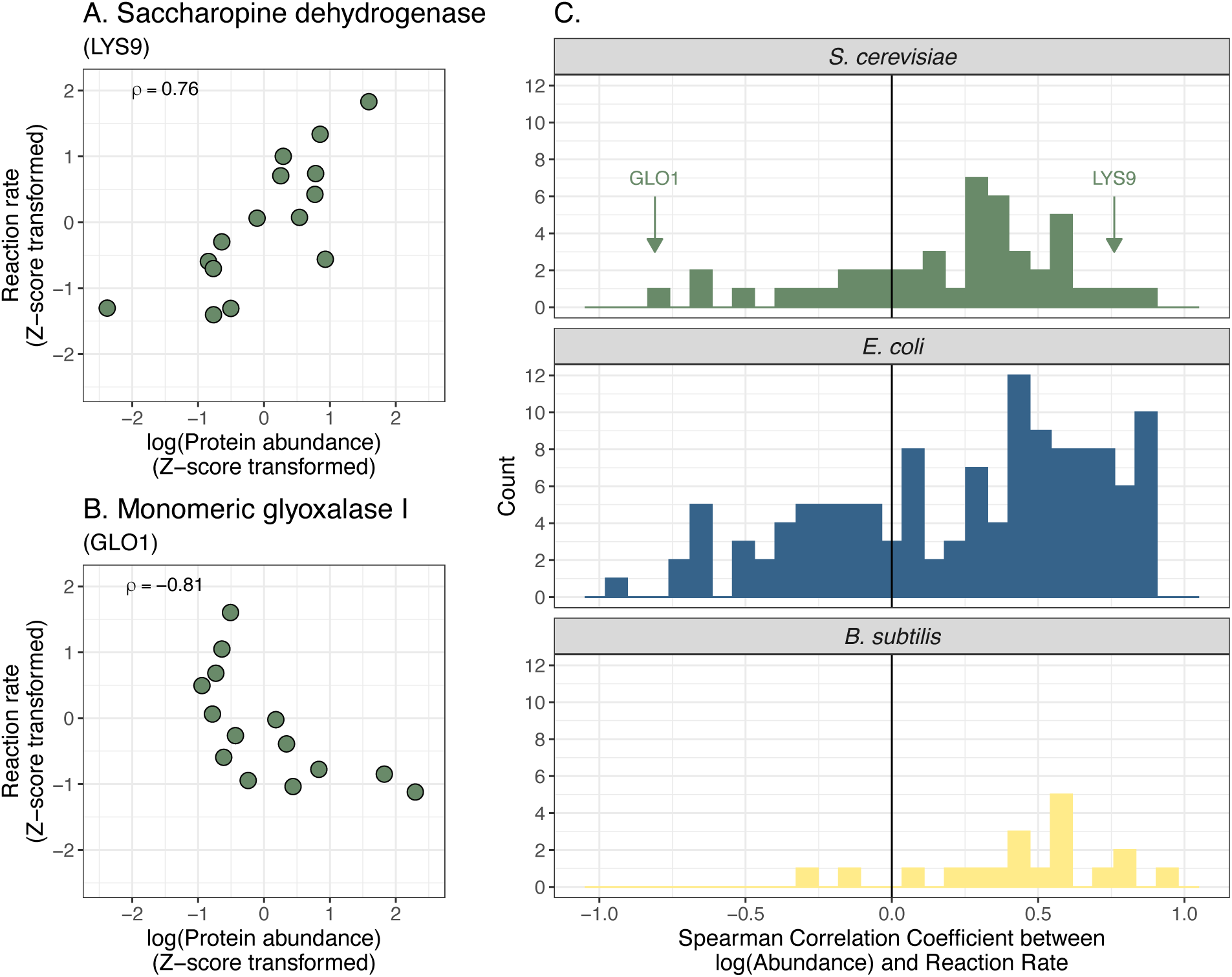
Empirical relationships between enzyme abundances and reaction rates. (A) The relationship between saccharopine dehydrogenase (LYS9) abundance and its associated reaction rate in *S. cerevisiae*. Protein abundances and reaction rates were log_2_-transformed prior to z-score transformation. ρ: Spearman correlation coefficient. (B) The relationship between monomeric glyoxylase I abundance and its associated reaction rate in *S. cerevisiae*. (A – C). (C) Distributions of protein-to-rate Spearman correlation coefficients for *S. cerevisiae, E. coli*, and *B. subtilis*.

Across all single protein-rate relationships in this dataset, most pairs exhibit weak covariation between protein abundance and rates (Fig. 2C). The median protein-rate relationship has a Spearman correlation coefficient of 0.25. The same trend holds when assessed with other functional forms of covariation, such as Pearson correlation with and without logarithmic transformation, mutual information, and a Bayesian hierarchical model to infer correlation coefficients (refs. 20, 21; Supplementary Fig. S1). To assess whether the limited covariation is due to measurement error, we examined protein abundance measurements between two closely related growth conditions and found them to be highly correlated (Pearson and Spearman correlation coefficients = 0.96, and 0.97, respectively; Supplementary Fig. S2), suggesting reproducible quantification. Derived reaction rates also had minimal measurement error (*R^2^* = 0.93 across independent studies, Fig. S3 in ref. 18). Together, these results show that enzyme abundance is typically not a good predictor of its corresponding reaction rate in budding yeast.

To expand this analysis to other taxa, we used previously published data from two bacterial species, *Escherichia coli* and *Bacillus subtilis* (22–26). Across a diverse set of steady-state growth conditions (25 and 8 conditions, respectively), reaction rates were derived alongside proteomic (*E. coli*) or transcriptomic (*B. subtilis*) measurements (see Materials and Methods). Similar to the budding yeast, covariation between rates and enzyme abundance spans a large range but is typically weak (Spearman correlation coefficients ranging from 0.93 to −0.94, with median = 0.24 and 0.45 for *E. coli* and *B. subtilis*, respectively).

In summary, we found highly variable and typically weak correlations between single enzymes and their cognate reaction rates in separate studies of three microbial organisms. We were unable to identify any features (e.g., ΔG of reaction or protein sequence similarity) that distinguish reactions with high versus low correlation coefficients (see Supplementary Information, Supplementary Figs. S3, S4). These results are also supported by other work that suggests metabolite concentrations have a large influence on reaction rates (e.g., refs. 6, 27, 28).

### Rate estimation using within-pathway proteins

We now turn the reaction rate estimation problem into a prediction task by leveraging additional proteins. We specifically examined whether reaction rates can be predicted by including other proteins within the same pathway. Conceptually, this question is equivalent to predicting rate using both enzymes in the branched pathway model (Fig. 1B). We used a statistical model in which the reaction rate is a function of within-pathway protein abundances (linear combination of log-transformed protein abundances). Similar to the single-enzyme analysis in the previous section, we used log-transformed protein abundances because protein abundances are often log-normally distributed. The model coefficients were determined using sparse regression methods, which prevent overfitting when there are many more predictors than observations. We used two-stage cross-validation to evaluate model performance.

Using the rate of *de novo* glutathione synthesis as an example, we demonstrate marked improvement in predictability when considering within-pathway proteins. Glutathione is one of the most abundant metabolites in yeast (29)and is a major player in redox metabolism and diverse ecological processes (28, 30–32). Dozens of proteins have been designated to glutathione metabolism, which consists of three parts: 1) *de novo* biosynthesis, 2) redox balance, and 3) degradation and transfer (Fig. 3A). We focused on the last step of *de novo* biosynthesis, which is catalysed by glutathione synthetase (GSH2) in *S. cerevisiae* (Fig. 3A). There was only a moderate relationship between GSH2 level and the rate of *de novo* glutathione biosynthesis (Spearman correlation coefficient = 0.5, *ρ^2^ =* 0.25; Fig. 3B), which makes GSH2 alone a poor candidate for this prediction task (Supplementary Fig. S5). By contrast, our statistical model (using ridge regression) trained on all quantified proteins in glutathione metabolism (15 proteins across 25 conditions, including 2 additional dilution series for auxotrophic strains) showed marked improvement (cross-validated *R^2^* = 0.585, Fig. 3C).

**Figure 3.**
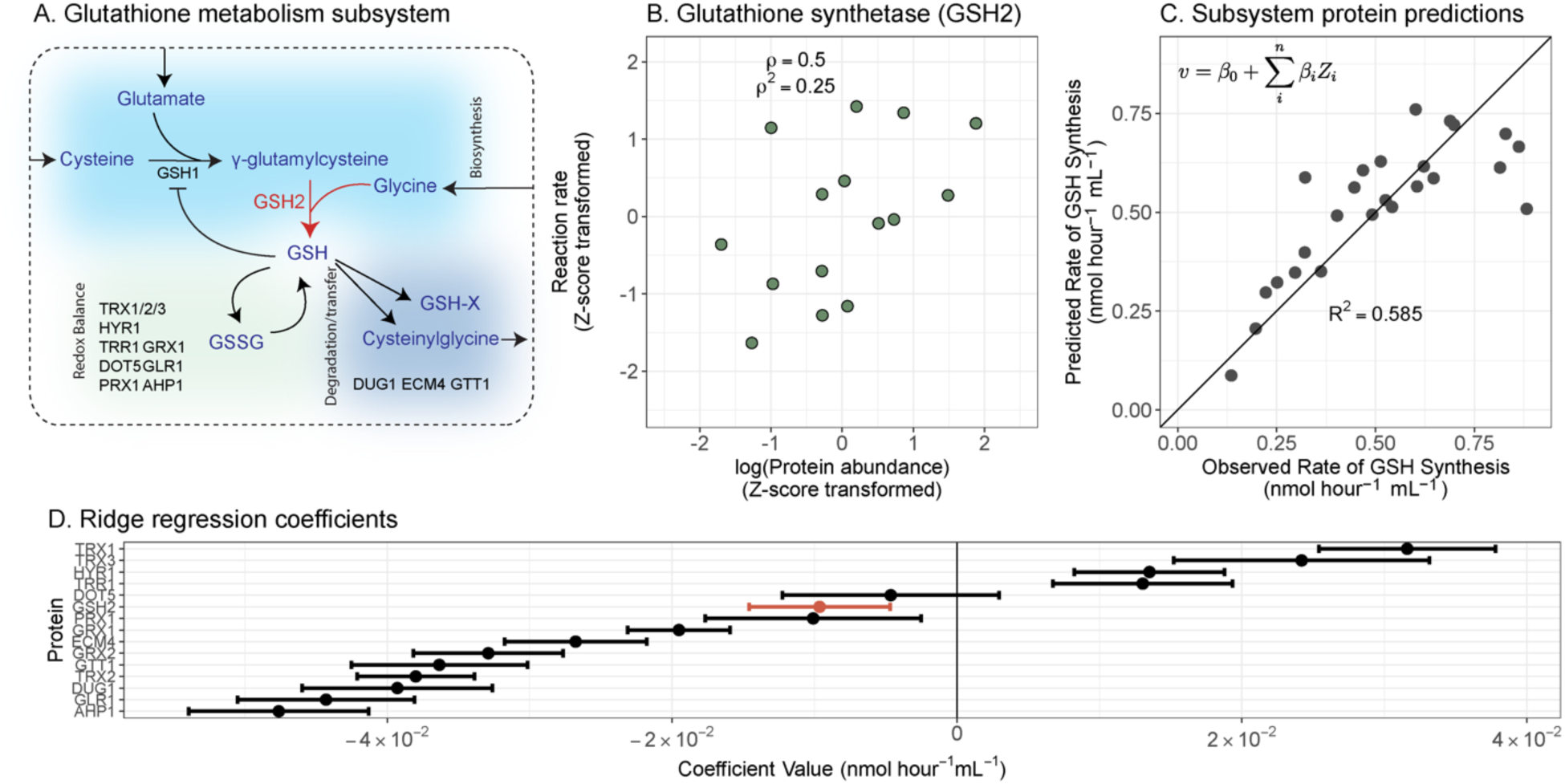
Estimating the rate of *de novo* glutathione biosynthesis using proteins within glutathione metabolism. (A) An overview of the structure of glutathione metabolism in *S. cerevisiae*, where individual proteins that were quantified (coloured black or red; except GSH1 which was not quantified but shown for context) are indicated in differently coloured sections according to their respective enzymatic role. GSH = reduced glutathione; GSSG = oxidised glutathione; GSH-X = glutathione bound to a xenobiotic. (B) The relationship between GSH2 protein abundance (log_2_-transformed and then z-score normalised), which mediates the *de novo* biosynthesis of glutathione, and its corresponding reaction rate (z-score transformed). ρ: Spearman correlation coefficient. ρ^2^: the coefficient of determination using ranked values. (C) Predicting the rate of *de novo* glutathione biosynthesis using other proteins within the glutathione metabolism subsystem. Each point is the mean of the predicted rate when it was left out of the training set. The cross-validated *R^2^* is displayed from leave-two-out cross-validation with protein abundances also z-score normalised to compare with panel. Ridge regression was used, and displayed are the resulting leave-two-out cross-validated predictions. The inset equation in panel C shows the general model structure used, where a reaction rate *v* is predicted as a function of coefficients (*β*_0_) and z-score transformed protein abundances (*Z_i_*). (D) All model coefficients (except for intercept coefficient) for a ridge regression model to predict glutathione synthesis. Error bars represent the standard deviation across cross-validation iterations, with 100 cross-validation iterations.

Notably, there was not a single dominant predictor protein in the pathway-level model. To compare the magnitudes of coefficients, we z-score transformed the log-transformed protein abundances. Several proteins have similarly high magnitude coefficients, indicating that they are collectively informative for the rate of *de novo* glutathione biosynthesis (Fig. 3D). Previous studies suggest that this rate is primarily controlled by the upstream enzyme, gamma-glutamylcysteine synthetase (GSH1; Fig. 3A; 29, 30, 33). However, GSH1 was not quantified in this dataset, suggesting that other proteins are also informative collectively. Several of the high-magnitude predictor proteins in our model (e.g., TRX1 or AHP1) were those balancing the redox state of glutathione. Reduced glutathione regulates its *de novo* biosynthesis by both competing with glutamate in the active site of GSH1 and influencing GSH1 expression (Fig. 3A; refs. (34–36). We speculate that these proteins are predictive for glutathione biosynthesis due to their roles in modulating the concentration of active GSH1.

More broadly, we found that many reaction rates in budding yeast are better predicted using within-pathway proteins (Fig. 4A-C). To define within-pathway proteins, we used the “subsystem” designations from a metabolic model based on KEGG annotations (19, 37, 38). Among 208 reactions that have >5 other proteins in the respective subsystem, more than half of had a cross-validated *R^2^* > 0.4 (Fig. 4C). By contrast, analogous ridge regression models trained solely on the focal enzymes perform much worse, with many cross-validated *R^2^ < 0*, indicating a prediction worse than a mean estimate (Fig. 4A, B). Note that the cross-validated *R^2^* (also called coefficient of determination) can be negative when the total variance is less than the residual variance based on the test set. Similar outcomes were observed for *E. coli* and *B. subtilis* datasets (Supplementary Fig. S6).

**Figure 4.**
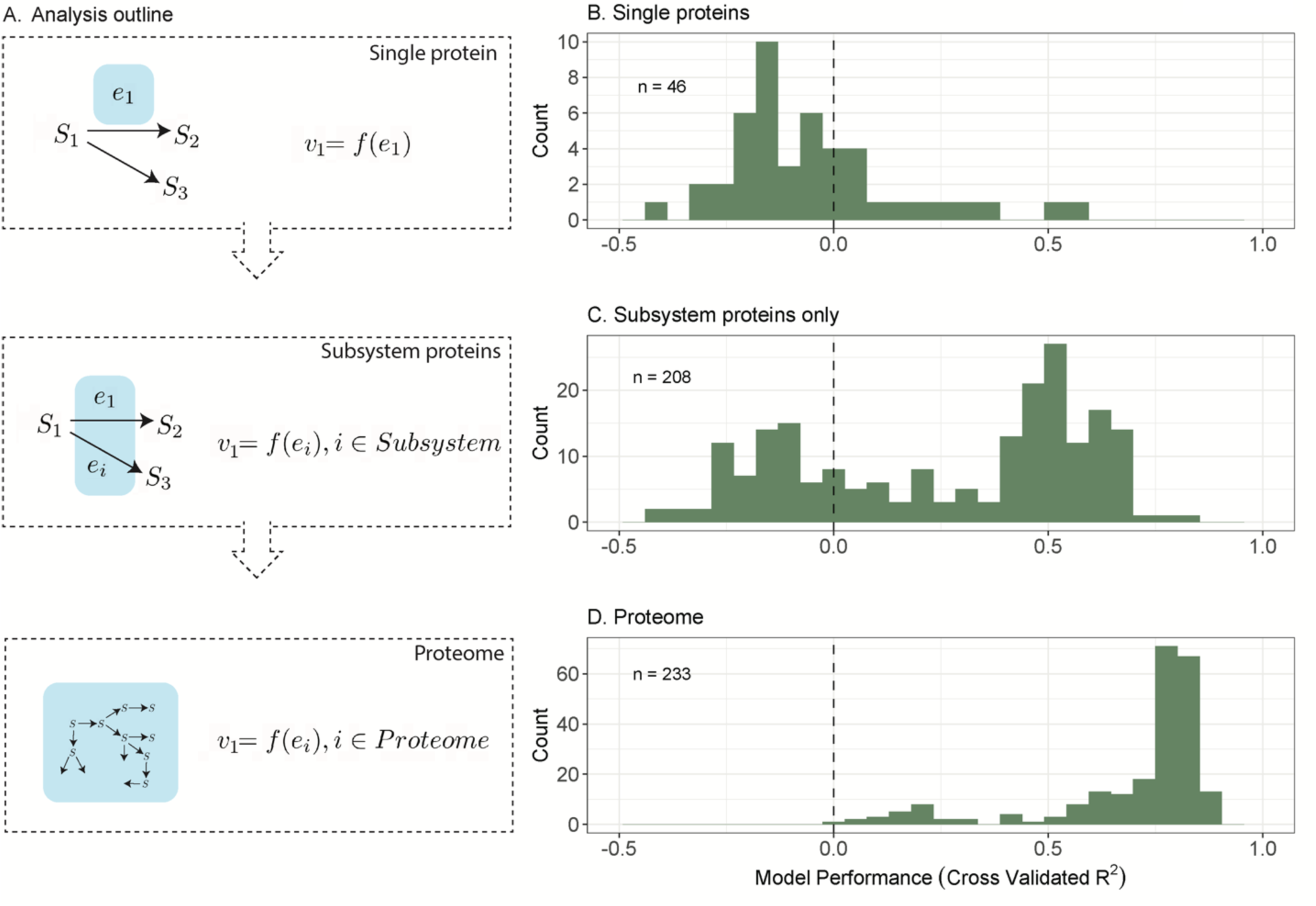
Predicting reaction rates using proteins at different scales. (A) Conceptual outline of the analysis approach. We used single proteins to predict their associated rates, and then used within-subsystem proteins, and then finally used the entire observed proteome. (B-D) Using proteomic data from *S. cerevisiae* (18), we predicted reaction rates and assessed model performance with the distribution of cross-validated coefficients of determination (*R^2^*). (B) Distribution of cross-validated *R^2^* using single proteins to predict rates. (C) Distribution of cross-validated *R^2^* using within-subsystem proteins to predict rates. (D) Distribution of cross-validated *R^2^* using within-subsystem proteins to predict rates. (B-D) Histograms are composed of different modelled rate predictions. Note that cross-validated *R^2^*< 0 corresponds to a prediction worse than a mean estimate (indicated by a value less than the dotted line). Single protein-based predictions were restricted to protein-to-rate pairs whereby 1) proteins had a single unambiguous reaction and 2) reactions had a single unambiguous protein, subsystem-level predictions only included available derived rates with more than five proteins observed per subsystem, and proteome-level predictions included all available derived rates.

From a practical perspective, these results suggest that proteins near a focal reaction rate may be a good starting point for improving rate estimates. For example, analyses of carbon fixation in phototrophs suggest that both RuBisCO and sedoheptulose bisphosphatase play important roles impacting carbon fixation rate (39), and therefore may be useful predictors for estimating carbon fixation rate. However, our results showed that within-pathway predictions have only moderate performance, and many rates could not be predicted using this approach.

### Rate estimation using proteomes

We expanded the regression analyses to include the proteome and discovered that many intracellular rates become highly predictable. Instead of relying on only proteins in the same subsystem, we now include all quantified proteins as predictors (1187 proteins). This additional information substantially increased model performance (233 rates) to a median cross-validated *R^2^* = 0.78 (compared to median *R^2^* = 0.4 for subsystems, Fig. 4A-D). Notably, 90% of predictive models for rates had a cross-validated *R^2^* between 0.39 and 0.87 (Fig. 4D). This finding was robust to subsampling the set of reactions to match single-protein predictions (46 rates, Supplementary Fig. S7), as well as to different cross-validation approaches (increasing the size of cross-validation test set, Supplementary Fig. S8; leaving whole media types out, Supplementary Fig. S9; or leaving out different growth rates, Supplementary Fig. S10). Of the various forms of cross-validation, leaving out media types had the largest impact on rate predictions with a median cross-validated *R^2^* of 0.66 (Supplementary Fig. S9). Together, these results suggest that proteome-wide abundances can be leveraged to predict reaction rates.

Contrary to the trend, there is a small subset of reaction rates that has lower model performance with a cross-validated *R^2^* of around 0.2 (Fig. 4D), which consisted mostly of reactions within the citric acid cycle. This group of rates were particularly sensitive to media-type leave-out cross-validation. We speculate that the statistical model would be improved if the training data included additional media types.

Interestingly, the predictions are good despite only requiring a small number of proteins. When using all quantified proteins to predict rates in 25 conditions, ridge regression suppresses the contribution of most proteins to avoid overfitting. To quantify the number of proteins that collectively make good predictions, we trained the same model using LASSO regression, which shrinks many fitting parameters exactly to zero (as opposed to asymptotically to zero in ridge). LASSO regression performed moderately well, albeit slightly worse, with a median cross-validated *R^2^* of 0.56 (Supplementary Fig. S11). We found that the number of proteins with non-zero LASSO coefficients (“predictor proteins”) ranged from 13-18 (25^th^–75^th^ quantiles; Supplementary Fig. S12). Therefore, only a small proportion of a proteome is required for accurately estimating reaction rates, which might be due to the highly modular gene regulatory network in *S. cerevisiae* (40).

Although the set of predictor proteins is different for different reactions, they are slightly enriched in translation machinery (Supplementary Materials and Methods), suggesting a potential connection between reaction rates and cellular growth rate. Indeed, growth is the emergent outcome of a complex network of reactions. We therefore determined if we were able to predict rates that are uncorrelated with growth rate. Succinate transport, for example, was uncorrelated with growth rate (Pearson correlation coefficient = 0.03), but the proteome-wide statistical model still performed well, with a cross-validated *R*^2^ = 0.78 (Supplementary Fig. S14). This result suggests that reaction rates do not need to be directly correlated with growth to be predicted. Nevertheless, we suspect that growth rate estimation using additional data types (e.g., genomic data; 39, 40) or gene expression data (43, 44) could help improve reaction rate estimation. We also emphasise that our current model ignores any explicit environmental context, and therefore environmental data could further improve predictions.

Overall, our results suggest that proteomic profiles contain information to accurately predict individual rates. We speculate that a similar result could be achieved with transcriptomic data.

## Discussion

Microbial reaction rates remain one of the most challenging observables in the environmental and biomedical sciences. Here we demonstrate that individual reaction rates only weakly correlate with their cognate enzyme abundance (e.g., Fig. 5A), but interestingly, accurate rate predictions are possible using statistical models trained on proteome-wide data (Fig. 5B). We anticipate that the increasingly accessible measurements for global gene expression will accelerate microbial rate estimation in natural environments.

**Figure 5.**
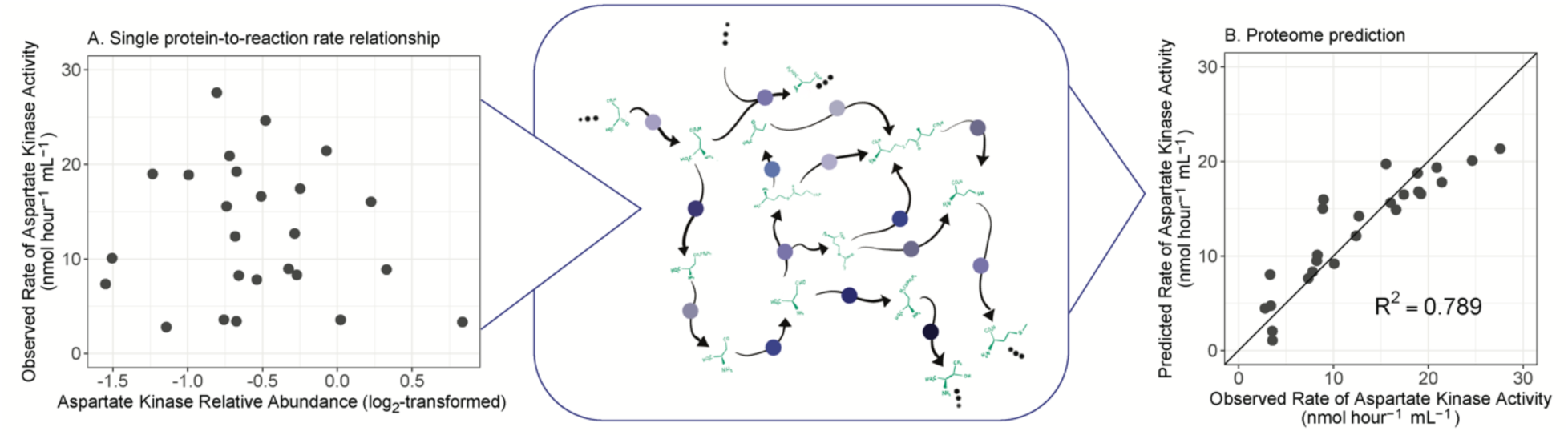
Proteomic context of reaction rate prediction. (A) Relationship between aspartate kinase relative abundance and aspartate kinase activity in *S. cerevisiae* across a range of culture conditions. This reaction is occurring in a larger network: the aspartate metabolism subsystem (center panel). Blue dots represent proteins mediating transformations of metabolites. (B) Reaction rate predictions when leveraging the entire proteomic context, which considers this focal reaction within the context of the broader network. The solid line plotted in panel B is the 1:1 line.

The lack of correlation between reaction rate and cognate enzyme abundance is consistent with evidence from biochemical considerations (Box 1), post-translational regulation (17), and predictions of the metabolic control analysis (45). However, this result defies a common implicit assumption used when interpreting gene expression data: an increase in enzyme abundance, say of ATP synthase, is an indicator for an increase in its overall activity, e.g., ATP production rate. The failed justification for this assumption has important consequences for interpreting differential gene expression across environmental settings.

At the pathway and proteome levels, we found that many rates become highly predictable given sufficient proteomic context (Fig. 5B), even without explicit mechanistic prior knowledge. One potential reason for this improved prediction is the distributed control points of reaction rates. The field of metabolic control analysis has suggested that, in many pathways, reaction rates are not controlled by a single, “rate-limiting” enzyme, but by several enzymes (45, 46). This conclusion is also evident from the branched pathway model presented earlier (Fig. 1B), where both *e_1_* and *e_2_* can influence the focal reaction rate *v_1_*. Distributed reaction rate control would enable prediction because additional proteins contain information about a focal reaction rate, which can then be learned via a statistical model.

However, it would be difficult to use our statistical models to infer pathway control points because protein abundances may merely correlate with, and not influence, pathway activity. In that case, the proteome is merely a reflection of distinct environmental context. Regardless of the mechanistic basis for the predictability, our finding indicates that proteomes encode information about reaction rates, and are sometimes sparse encoders, requiring only a small number of proteins.

We next highlight five important considerations for extending our approaches to natural microbial assemblages. First, we predicted reaction rates in steady state cultures. An important next step is to determine whether our results can be generalized to cultures that are not in steady-state growth. Natural environments will present additional challenges for these approaches, and future work should consider how environmental dynamics impact the ability of these methods to accurately estimate reaction rates.

Second, we estimated reaction rates using proteomes from a single species. It is unclear whether a model trained on a single species would be capable of accurately estimating rates of another taxon, which is a major hurdle when applied to natural communities. Methods from machine learning, in particular transfer learning, which allows predictions to be extended across biological systems, hold promise in tackling this problem (47).

Third, we examined biomass-normalised rates (mmol ⋅ hour^-1^ ⋅ cell volume^-1^), which may not always be the most informative normalization. If a given rate is proportional to biomass, therefore leading to constant biomass-normalised rates, the rate prediction might perform poorly (i.e., a low *R*^2^). However, in this scenario, estimating biomass alone could inform rates in natural environments. For example, if the rate of nitrogen fixation *per nitrogen fixer biomass* only varied ∼20% around the mean, then determining nitrogen fixer biomass alone may suffice. Determining taxon-specific biomass is still a non-trivial problem given variation in the ratio of genomes, transcriptomes, and proteomes to total taxon-specific biomass (48).

Fourth, we were able to predict reaction rate in some instances using only within-pathway proteins. This result suggests that targeted proteomic methods used for estimating *in situ* rates might look to proteins nearby within a given pathway for improving rate estimates. For example, proteins from within amino acid biosynthesis subsystems could improve estimates of glutamate biosynthesis rates.

Finally, the subjective quality of rate estimates is ultimately a function of how they are used, and these types of estimates can be used in many different settings. For example, estimates of toxin production rate by a pathogenic bacteria may be very uncertain, but still informative.

Overall, we have proposed approaches for leveraging the vast amounts of molecular data to make quantitative estimates of microbial processes, and our demonstration using cultured organisms showed that proteome-level information has promise for estimating *in situ* rates. Converting the units – i.e. from proteomic composition to reaction rates – is a step towards bridging molecular data with ecosystem-scale models.

## Materials and Methods

### Datasets descriptions

The *S. cerevisiae* dataset (18) consists of five media types used to limit growth in chemostats (glucose-limited, ammonia-limited, phosphorous-limited, leucine-limited, and uracil-limited), each with 5 different dilution rates. In the *E. coli* dataset, there were a wide range of culture conditions aggregated (see ref. 39–42). The *B. subtilis* dataset contained 8 different conditions that differed based on the carbon source (e.g., glucose, fructose, gluconate, etc.; ref. 40).

Reaction rates were derived in different ways depending on the dataset. Reaction rates were derived in *S. cerevisiae* as described in the main text. For *E. coli*, rates were derived using parsimonious Flux Balance Analysis (4, 39–42; in units of mmol ⋅ hour^-1^ ⋅ gCDW^-1^; gCDW: gram of cell dry weight) that leveraged associated growth rates. For the *B. subtilis* dataset, rates were inferred for an isotopically-labelled culture using an isotopomer balancing model constrained by the substrate uptake and product release rates (refs. 25, 26, with units of mmol ⋅hour^-1^ ⋅ gCDW^-1^).

Protein or transcript abundances were quantified differently for each dataset (*S. cerevisiae* proteomic quantification described in main text). For *E. coli*, protein abundance was quantified using mass spectrometry and is in units of mmol gCDW^-1^ (see details for unit conversions in ref. 39). For *B. subtilis*, transcripts were quantified using a microarray and are in units of relative abundance (which is reflective of molar abundance and relative to total mRNA).

### Analyses of single protein-to-rate relationships

For our analyses of covariation between single proteins and their associated rates, we only considered model-derived reactions that have a single corresponding protein (and these proteins also have only a single corresponding reaction). This criterion resulted in 46 and 125 protein-to-rate relationships for *S. cerevisiae* and *E. coli,* respectively, and 19 transcript-to-rate relationships for *B. subtilis*. Note that for the *S. cerevisiae* dataset, leucine- and uracil-limited growth had genetically modified auxotrophs. We therefore excluded these 10 experimental conditions for the first analysis on single protein-to-rate relationships, because the genetic modifications themselves may have driven the observed relationships. All code to reproduce analyses and figures are available online at https://github.com/jspmccain/pred-env-ge.

### Predicting rates using single proteins, pathways, and whole proteomes

We predicted microbial reaction rates using various types of regression models. Protein abundances were log_2_-transformed prior to fitting these regression models. For examining coefficient magnitudes in the within-pathway predictive models, we additionally z-score transformed protein abundances across different conditions (after log_2_-transformation). The statistical model we used to predict the rates is:

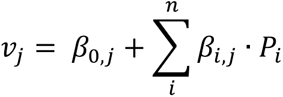

where *v_j_* is the *j*th reaction rate, *β* are the model coefficients, and *P* is the log_2_-transformed protein abundances (or additionally z-score transformed for Fig 3C, D). The summation is over *n* proteins, and each protein has a corresponding coefficient (subscript *i*). The value of *n* varies depending on whether single proteins, within-subsystem proteins, or proteome-wide models are used. The model coefficients were estimated using either ordinary least squares (for single proteins), ridge regression, or LASSO (“Least Absolute Shrinkage and Selection Operator”) regression.

Simple linear regression was used for evaluating the predictive ability of a single protein for a single rate. For evaluating single protein predictions in the *S. cerevisiae* and *E. coli* datasets, we used leave-two-out cross-validation. For the *B. subtilis* dataset, we only looked at rates that had at least 5 rate values across culture conditions (leaving 10 reactions that have at least 5 rate values), and we also only used leave-one-out cross-validation for this dataset given the small size. Cross-validation is required, as opposed to transforming correlation coefficients to compare predictions across different models. We analysed all rate-to-protein pairs shown in Fig. 2C for the *E. coli* and *S. cerevisiae* datasets.

For within-subsystem protein predictions, we restricted our analyses to the *S. cerevisiae* dataset for two reasons. 1) There is too little data in the *B. subtilis* dataset to assess predictive models with many predictor variables (8 conditions; given the cross-validation scheme described above). 2) The set of inferred reaction rates in the *E. coli* dataset were highly correlated and therefore we cannot independently assess predictions across rates (most derived rates had a Pearson correlation coefficient > 0.99 with each other). Note that in the *S. cerevisiae* dataset, there were two media types in which genetically modified auxotrophs were growing. We excluded them for the first analysis on single protein-to-rate relationships, but we included them in the analyses using statistical regression models.

We predicted individual rates by first considering only proteins from within a given pathway. For example, we predicted the rate of *de novo* glutathione biosynthesis using proteins designated within the “Glutathione Metabolism” subsystem (subsystems are mostly based on KEGG Pathways, 27, 28), we specifically follow subsystem classifications described in the yeast-GEM version 8.6.3 model (19). We restricted our analysis to pathways with more than five quantified proteins per subsystem (19), which was sufficient for the two-stage cross-validation approach described above.

For evaluating model performance for both within-pathway and proteome-wide predictions, we used several different types of cross-validation and then computed the coefficient of determination using the test set (*R^2^*). Specifically, we used: 1) leave-two-out cross-validation, 2) leave-five-out cross-validation, 3) cross-validation by leaving out unique growth rates, and 4) cross-validation by leaving out limiting substrate type. For the first two cross-validation approaches, we left two or five experimental conditions out of the training, and trained a model on the remaining conditions, and then assessed remaining test set predictions (using 100 cross-validation iterations). When a LASSO or ridge regression model is trained on the remaining conditions, the penalty parameter (*λ*) is determined with ∼10-fold cross-validation using the R package glmnet (51). Cross-validation for limiting substrate or growth rate was done by removing the five growth rates all belonging to the same media type (e.g., all the ammonia-controlled growth conditions), or the same growth rate belonging to different media types.

We calculated the cross-validated *R^2^* as follows: *R^2^ =* 1 *-SSE/SST*; where *SSE* is the sum of squared errors using only the out-of-training set predictions, and *SST* is the total sum of squares (using the observations). Note that some reactions have zero rates except for one condition (uracil phosphoribosyltransfer, L-leucine transport, and uracil transport). In these cases, the cross-validation with limiting substrate or leave-5-out cross-validation cannot be used to train predictive models due to constant response values.

## Acknowledgements

This work was aided by discussions with Daniel Segré, Rogier Braakman, and people from the Simons Collaboration on Computational Biogeochemical Modelling of Marine Ecosystems. Thank you to Robert Battaglia for comments on the manuscript. J.S.P.M. is a Damon Runyon Fellow supported by the Damon Runyon Cancer Research Foundation (DRG-2470-22). GLB was supported by the Simons Foundation (award # 1195558). MJF is grateful for support from the Simons Foundation (CBIOMES grant 549931 to MJF). This work is supported by NIH 5R35GM124732.

