## Supplemental Materials and Methods for "Microbial reaction rate estimation using proteins and proteomes"

**Supplementary Materials for "Microbial reaction rate estimation using proteins and proteomes"**

### 1. Comparing estimated correlation coefficients for homologous proteins

We examined if homologous proteins (between *E. coli* and *S. cerevisiae*) have similar relationships between protein abundances and *in vivo* rates. We compared sequences using BLASTP (using default settings with version 2.6.0+) from *E. coli* to *S. cerevisiae*, and then matched proteins if they had an E-value cutoff of $<1 \times 10^{-10}$ (i.e., highly similar based on sequence). The relationship between the estimated correlation coefficients across taxa was weak (Supplementary Fig. S1).

### 2. Quantifying single protein-to-rate relationships using various functional forms

Across the three datasets described in the main text, we examined the Pearson correlation coefficient (with and without log-transformation of the abundance data), the Spearman correlation coefficient, and the mutual information (with log-transformation of the abundance data). Mutual information was calculated using B-spline functions (1, 2). Note that calculating mutual information with few observations, for example as in the *B. subtilis* dataset, is typically sensitive to the number of bins and therefore can be difficult to interpret. The number of bins used for calculating mutual information was set as the number of values per relationship to the power of ⅓.

**3. Quantifying single protein-to-rate relationships using a Bayesian regression model**

We fitted a Bayesian hierarchical regression to each of the individual datasets with the following structure. Protein abundances were $\log_2$-transformed, and then both reaction rates and protein abundances were *z*-score transformed. Z-score transformations were calculated by protein, where the standard deviation and mean for a given protein is calculated across conditions. Note that the slope of a linear regression, where the independent and dependent variables are z-score transformed, is equivalent to the Pearson correlation coefficient. Throughout, we used uniform priors, with standard deviations bounded by zero:

$$\gamma \sim U(-1, 1)$$

$$\tau \sim U(0, \infty)$$

$$\sigma \sim U(0, \infty)$$

$$\beta_0 \sim U(-100, 100)$$

$$\beta_1 \sim Normal(\gamma, \tau^2)$$

$$Rate_i^z \sim Normal(\beta_0 + \beta_1 \cdot Protein_i^z, \sigma^2)$$

Above, $\gamma$ is a parameter describing the mean of the individual slope estimates ($\beta_1$), $\tau$ is the standard deviation of this distribution of individual slope estimates, and $\sigma$ is the standard deviation of the group-level distribution of slopes. $Rate^z$ and $Protein^z$ refer to the z-score transformed reaction rate and protein abundance, respectively. Every protein-to-rate relationship is parameterized with a partially-pooled, individual, slope estimate ($\beta_1$), as well as an independently estimated intercept $\beta_0$. We constrained prior probability distribution of the slope estimates to be between -1 and 1, to compare with Pearson correlation coefficients, which are similarly constrained between -1 and 1 (although this constraint only slightly modified the highest estimated $\beta_1$ values' credible intervals). One advantage for using a Bayesian model is that uncertainties are represented as probabilities, and the hierarchical nature of this modelling approach allows information to be shared across coefficient estimates. All Bayesian hierarchical regressions were fitted using Hamiltonian Monte Carlo, written in Stan and Rstan (3, 4).

**4. Explaining variation in protein-to-rate relationships**

We also examined protein-level features that explain variation in the slope of the relationship between z-score transformed protein abundance (first $\log_2$-transformed) and z-score transformed reaction rate by extending our Bayesian hierarchical model. We explored three hypotheses for why a given protein has a higher slope coefficient in the normalised protein abundance to rate relationship: 1) proteins with a higher mass fraction have higher coefficients, 2) proteins with more connected substrates have higher coefficients, and 3) proteins mediating reactions with a lower $\Delta G$ have higher coefficients, First, we describe the rationale for each of these hypotheses, and then we describe how these hypotheses were tested in the Bayesian hierarchical regression.

1) Protein production is costly (5, 6), and we therefore hypothesise that more abundant proteins tend to be more highly correlated with their corresponding reaction rates. Protein abundance is, however, contingent on the set of conditions examined, so we averaged mass-weighted protein abundance across different conditions using the raw data from ref. (7). Specifically, we first normalised all peptide abundances using the sum of isotopically

light peptide intensities as a normalisation factor. This approach adjusts for varied amounts of protein injected across samples. We then summed up these normalised peptide intensities *per protein*, which is analogous to calculating a proteomic mass fraction, which assumes a low variability in peptide length across peptides. This approach is analogous to others that have leveraged the notion of proteomic mass fraction as a proxy for proteomic investment (8, 9).

2) We theoretically examined how substrate sharing across enzymes might influence the relationship between enzyme abundance and rates. Consider a single substrate, where the dynamics can be described by the balance between uptake ($\rho$), consumption by at least one enzyme ($E_i$, where $i$ refers to a specific enzyme), and dilution by growth ($\mu$). These processes can be represented by a single ordinary differential equation (where we rescaled $E_i$ by the turnover number for simplicity):

$$\frac{dS}{dt} = \rho_s - S \sum_{i}^{n} E_i - \mu S$$

Solving for the steady state levels of the substrate, *S*, we can then arrive at an expression for a rate mediated by a single enzyme (in the n = 1 case):

$$V_1 = E_1 \frac{\rho_S}{E_1 + \mu}$$

We can then solve for a system where *n* enzymes are consuming a single substrate *S*:

$$V_i = E_i \frac{\rho_S}{\sum_1^n E_n + \mu}$$

Assuming that the update rate is constant, this model predicts that as the number of proteins per substrate increases, there is additional variability introduced into the relationship between protein abundance and reaction rate. This model is a generalization of our branched pathway model in Box 1, such that *n* enzymes can consume a single substrate. Variation in the abundance of non-focal proteins can therefore alter the concentration of the single substrate, thus inducing variation in the reaction rate. We therefore calculated a metabolite-specific score: the number of shared proteins per metabolite. Each reaction mediated by a protein typically has multiple substrates. So, for each reaction, we then calculated the geometric mean of each of these metabolite specific scores. The geometric mean provides a score that is less sensitive to inclusion of extremely connected substrates, like ATP or GTP.

3) Classic work on reaction thermodynamics suggests that reactions that have a near-zero $\Delta G$ are less likely to be modulated by enzymes. This theory is because of the 'force-flux' relationship (10): reactions with near-zero $\Delta G$ values have, by definition, similar forward and reverse fluxes. Therefore, an increase in enzyme abundance will contribute not only to the (presumably) beneficial forward reaction, but also to the reverse reaction. This logic suggests that reactions with a very negative $\Delta G$ are more likely to be controlled by enzyme abundance (e.g., 6, 7). We therefore examine how much variation in the slope in between normalised protein abundance and normalised reaction rate varies as a function of $\Delta G$ (using previously published data on reaction thermodynamics, refs. 8, 9).

We used a Bayesian hierarchical model to explain variation in the slope of the relationship between z-score transformed enzyme abundance (first $\log_2$-transformed) and z-score transformed reaction rate in the *S. cerevisiae* and *E. coli* datasets, testing the hypotheses described above. (The *B. subtilis* dataset only included 19 transcript-to-rate pairs and therefore did not have sufficient data for these purposes). This model is an extension of the previous Bayesian hierarchical model, and so we jointly estimate both the linear relationship between protein abundance and *in vivo* rate (across different experimental conditions), and we estimate coefficients that explain variation in this slope (across different proteins).

$$\tau \sim U(0, \infty)$$

$$\sigma \sim U(0, \infty)$$

$$\gamma_0 \sim U(-\infty, \infty)$$

$$\gamma_1 \sim U(-\infty, \infty)$$

$$\gamma_2 \sim U(-\infty, \infty)$$

$$\gamma_3 \sim U(-\infty, \infty)$$

$$\beta_0 \sim U(-\infty, \infty)$$

$$\beta_1 \sim Normal(\gamma_0 + \gamma_1 \cdot Mean\ Protein\ Abundance \ldots$$

$$\ldots + \gamma_2 \cdot Substrate\ Connectivity\ Metric + \gamma_3 \cdot \Delta G\ of\ Reaction, \tau)$$

$$Rate_i^z \sim Normal(\beta_0 + \beta_1 \cdot Protein_i^z, \sigma^2)$$

where $\gamma_1$ is a coefficient corresponding to protein cost hypothesis, $\gamma_2$ is a coefficient corresponding to variation in enzyme substrate connectivity, and $\gamma_3$ is a coefficient corresponding to variation in $\Delta G$ of a reaction. We used a uniform prior on each of these coefficients, as well as the standard deviation parameters ($\tau$, and $\sigma$), except these latter parameters were bounded by zero.

**5. Gene Ontology Enrichment Analysis**

We examined which GO terms were enriched in the predictor proteins for the statistical models with good performance. Specifically, we examined whole proteome LASSO regression results with a cross-validated $R^2 > 0.5$ (using leave-two-out cross-validation). We then determined which GO terms are enriched in these models, compared to the set of GO terms associated with the observed proteins. A binomial distribution can describe the number of proteins associated with a single GO term and the number of proteins associated with all GO terms. Specifically, we modelled the number of proteins that are labelled with a given GO term as follows:

$$Y_i \sim Binomial(N_i, p_i),$$

where the number of proteins labelled with the $i$th GO term (e.g., "regulation of translation") follows a binomial distribution parameterized by $N_i$ (the total number proteins labelled with all GO terms), and the success probability parameter $p_i$ (the number of proteins labelled with the $i$th GO term). The null expectation is formulated as a similarly parameterized a binomial distribution, except using all proteins that were included in the cross-validated LASSO models (above our $R^2$ cutoff). $N_i$ is also the total number of proteins labelled with all GO terms (which is dependent on the total number of cross-validation iterations), and the success probability parameter $p_i$, the number of proteins labelled with the $i$th GO term from the LASSO models. After fitting these two binomial distributions, we compared the estimated $p_i$ from the LASSO models with those from the null model. We specifically examine the difference in their posterior probability distributions for the estimated parameters $p_i$ (see Supplementary Figures S13). GO terms were retrieved (February 17$^{th}$, 2024) from GO-Slim mapper from the *Saccharomyces cerevisiae* Genome Database website (https://www.yeastgenome.org/goSlimMapper). We specifically focused on the "Process" GO term category.

**6. Exploratory analyses of within-pathway predictive model coefficients**

Using the within-pathway sparse regression models, we explored whether there are certain proteins that are typically predictive of rates within pathways. Further, we examined if there were any unique characteristics among these proteins. For example, we evaluated if the position in a pathway relates to reaction rate predictive capacity (i.e., the magnitude of coefficients in ridge regression). To do so, we re-ran our ridge regression models with scaled protein abundances (*z*-score normalised) and examined pathways that had a median $R^2 > 0.5$ for all reaction rate prediction models (10 pathways in total, with 67 individual derived rates within these pathways). From these pathways, we hypothesized that proteins with high magnitude coefficients (either very low or very high) were proteins that mediated reactions with a low $\Delta G$ of reaction. We also examined if there was a relationship between network-based proximity of the predictor protein and the rate being predicted. For the network-based proximity metric, we specifically converted the metabolic network into a graph, where nodes of the graph are reactions, and edges are present when two reactions share a reactant or a product. We could not identify any relationship between this graph-based distance and the strength of the coefficients, within pathways, nor could we identify if the $\Delta G$ is associated with coefficient strength.

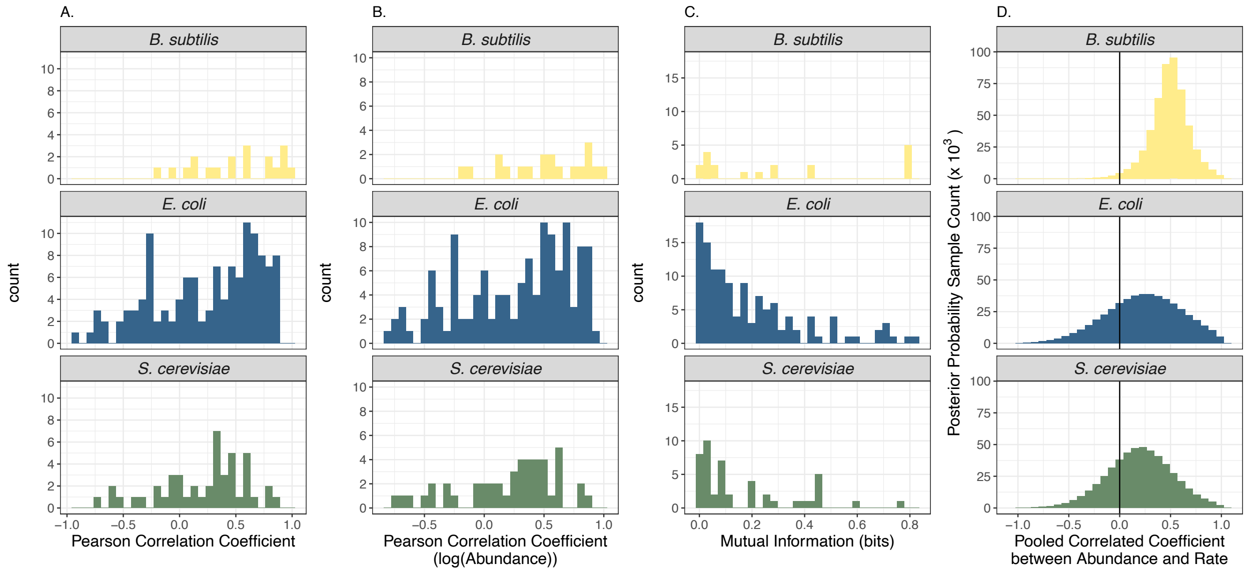


**Figure S1.** Distributions of various statistical relationships between protein or transcript abundance and corresponding reaction rates across three taxa. (A) Pearson correlation coefficients, without log transforming either rates or abundances, (B) Pearson correlation coefficients, with $\log_2$-transformed abundances only, (C) mutual information using raw values for both rates and abundances, and (D) pooled correlation coefficient using a Bayesian model. In the Bayesian model, the pooled correlation coefficient is the group-level distribution of slopes from the Bayesian hierarchical regression, where z-score normalised and $\log_2$-transformed protein or transcript abundances were regressed with z-score normalised reaction rates, and a linear model was fitted (see details in Supplementary Materials and Methods).

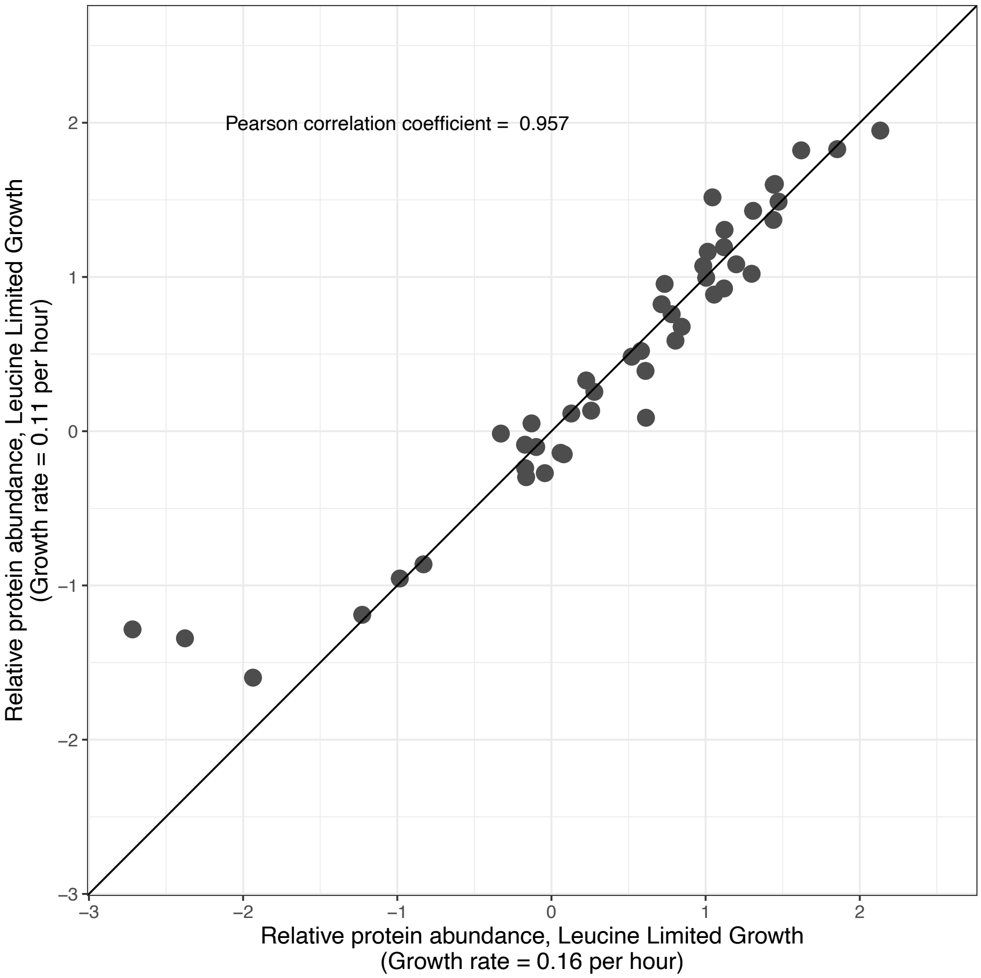


**Figure S2.** Correlation between the relative protein abundance across two similar conditions with different dilution rates in *S. cerevisiae*, data from ref. 1. Protein abundances were $\log_2$-transformed (46 proteins in total).

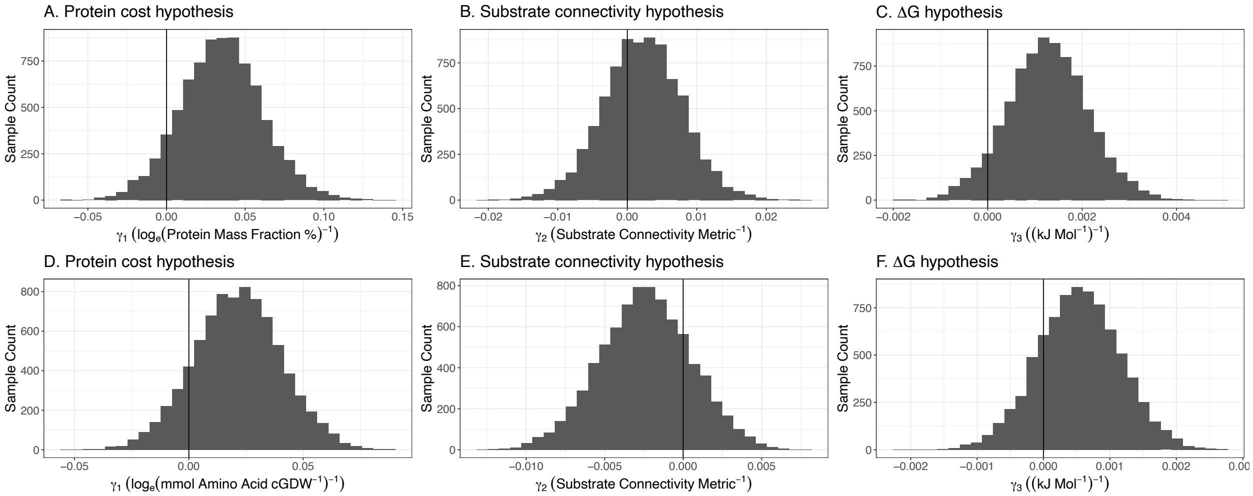


**Figure S3.** Posterior probability distribution samples for the Bayesian hierarchical model examining variation in slopes across *S. cerevisiae* (top, A-C) and *E. coli* (bottom, D-F). The first column quantifies the coefficient corresponding to protein mass fraction for *S. cerevisiae* and *E. coli*. The middle column (B, E) quantifies the coefficient for our substrate connectivity metric (see Supplementary Materials). The last column (C, F) shows the coefficient estimate for the ΔG of reaction. Coefficient estimates for panels A and D can be interpreted approximately as: a 1% increase in the protein mass fraction percentage (A) or mmol AA $gCDW^{-1}$ (D) is associated with a $\gamma_1/100$ increase in the Pearson correlation coefficient. Coefficient estimates for panels B, C, E, and F can be interpreted as the change in Pearson correlation coefficient as consequence of a 1 unit change in the dependent variable (e.g., $\Delta G$).

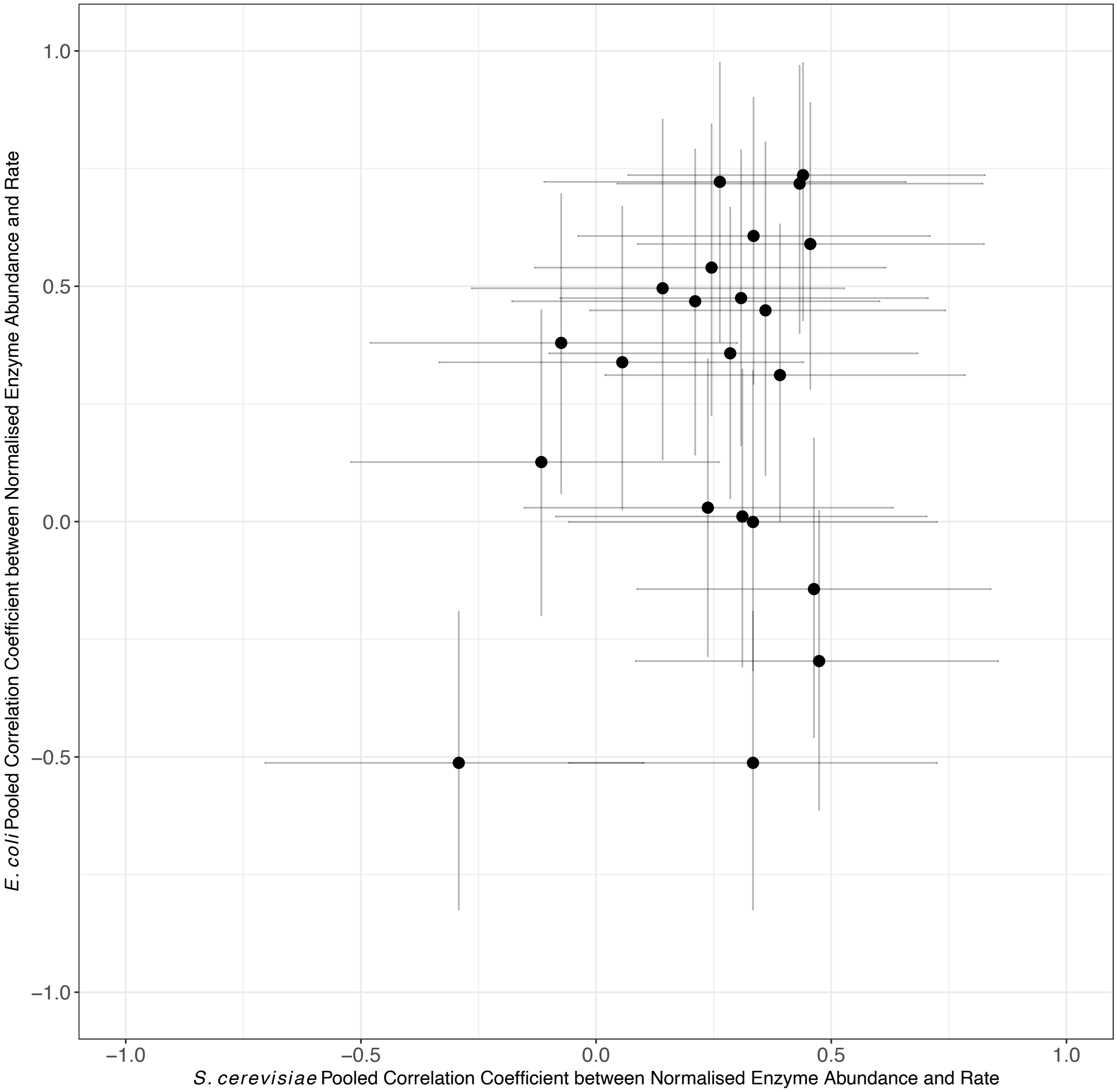


**Figure S4.** Comparison of pooled correlation coefficients in *S. cerevisiae* and *E. coli* between homologous proteins. Pooled correlation coefficients are the slope estimates from the linear regression between z-score and $log_2$-transformed enzyme abundances and z-score transformed rates (see Supplementary Materials and Methods). Using the median posterior estimates, the Pearson correlation coefficient is 0.24, with 22 protein pairs in total.

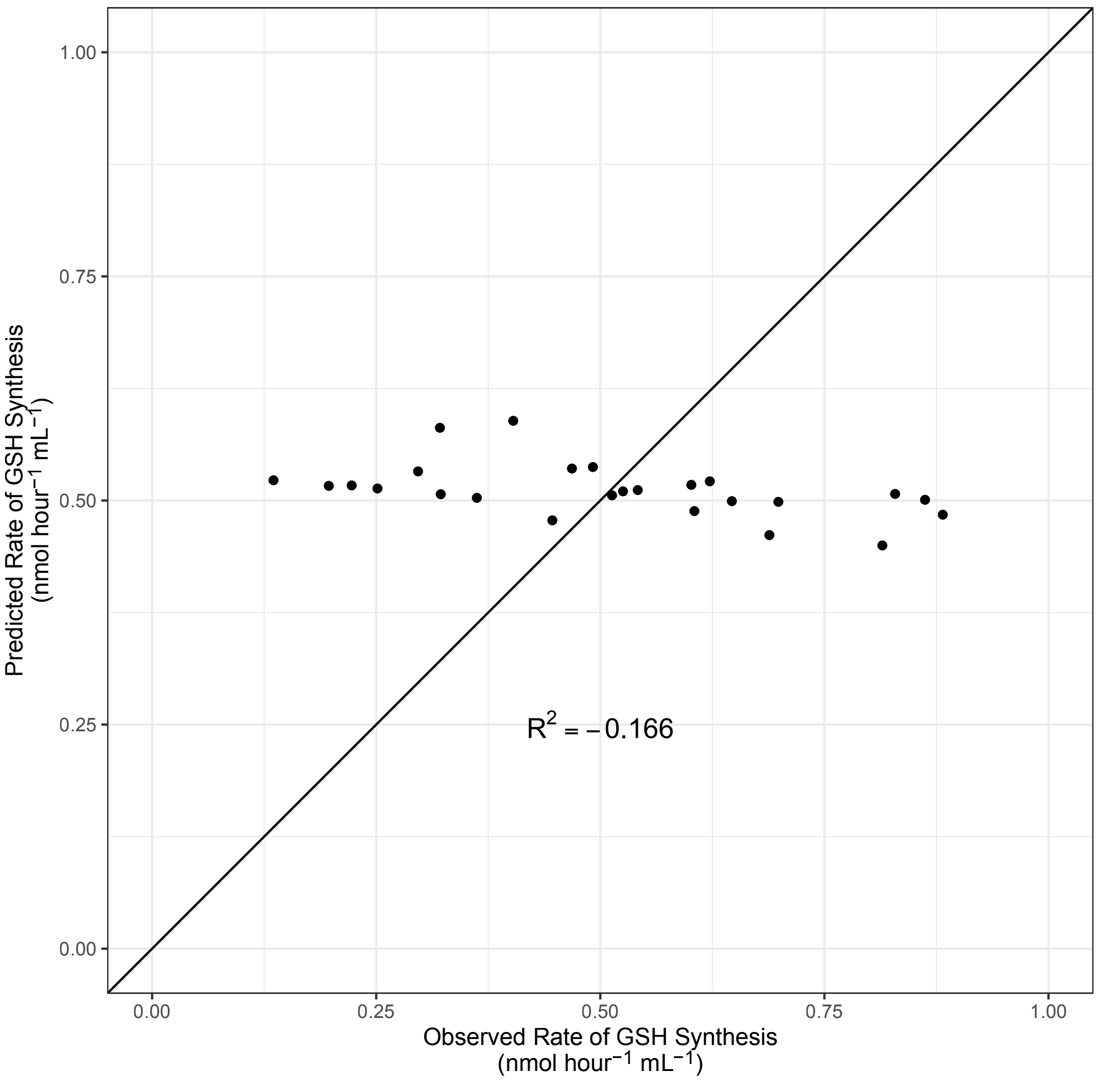


**Figure S5.** Predicting the rate of *de novo* glutathione biosynthesis as a function of z-score normalised and $log_2$-transformed GSH2 abundance, using a linear regression with leave-two-out cross-validation. Z-score normalisation was done to compare with Fig. 3C in the main text (this plot also includes 25 individual rate predictions). Cross-validated coefficient of determination ($R^2$) is shown, where a negative value indicates a model worse than a mean estimate.

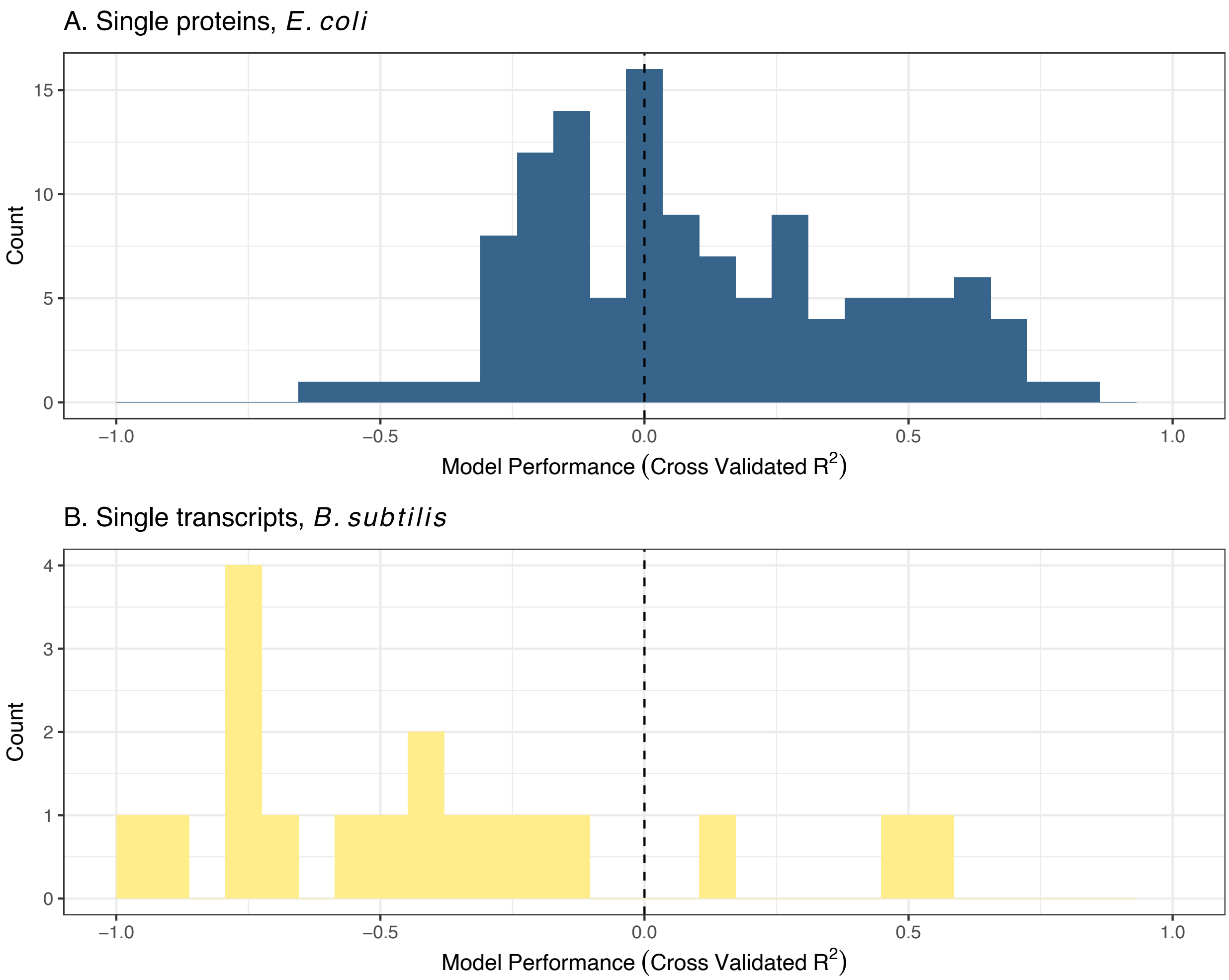


**Figure S6.** Linear regression model performances of reaction rate predictions using (A) single proteins and (B) single transcripts for the *E. coli* (125 rate predictions using single proteins) and *B. subtilis* (19 rate predictions using single proteins) datasets, respectively. Abundances were first $\log_2$-transformed, and model performance was assessed using cross-validated coefficient of determination ($R^2$). A negative value indicates a model worse than a mean estimate.

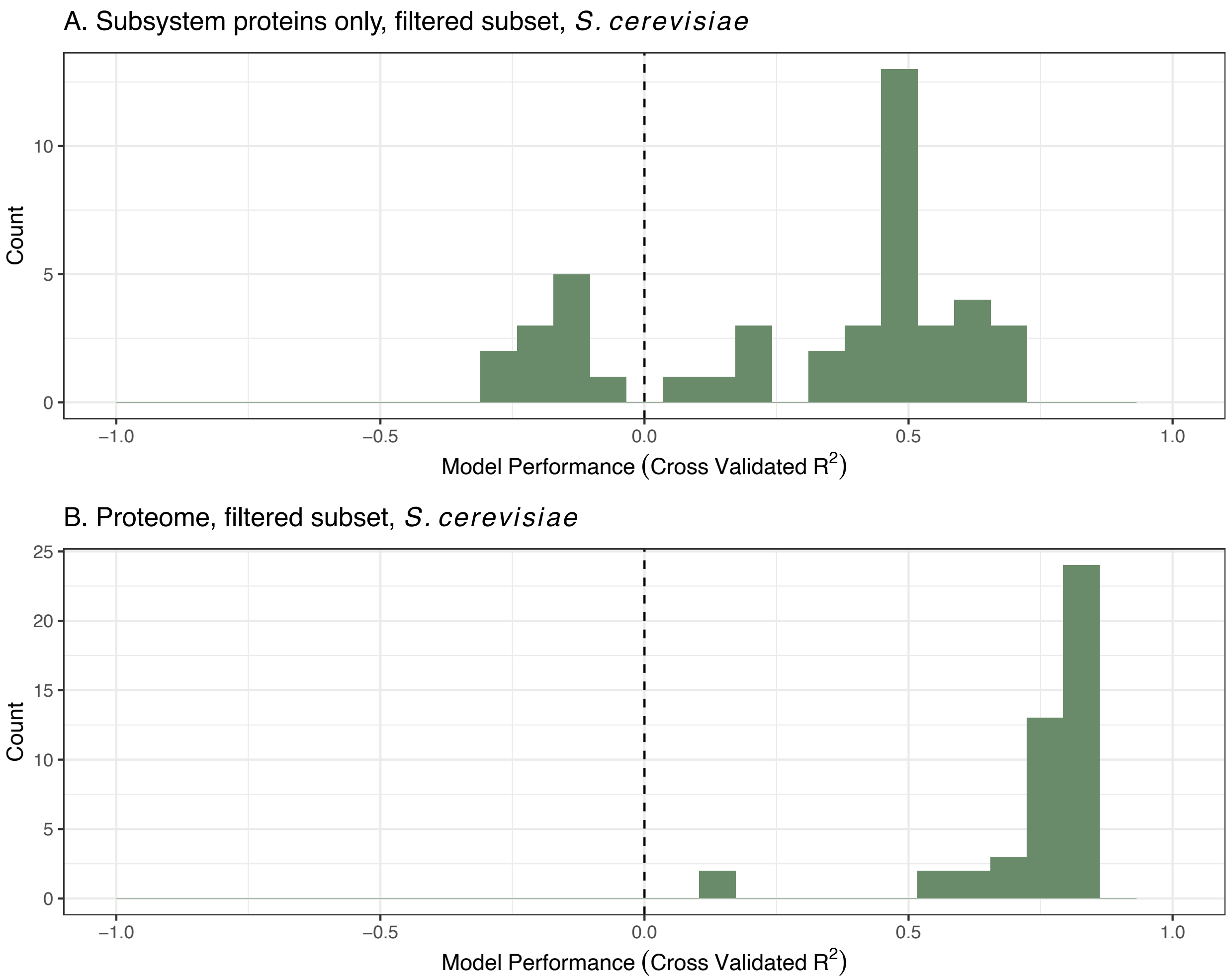


**Figure S7.** Distributions of cross-validated $R^2$ using leave-two-out cross-validation for both the pathway level (A) and proteome level predictions (B), using ridge regressions. In this case, only the same subset of proteins that were used for single proteins are shown (46 predictive model summaries in total in each panel).

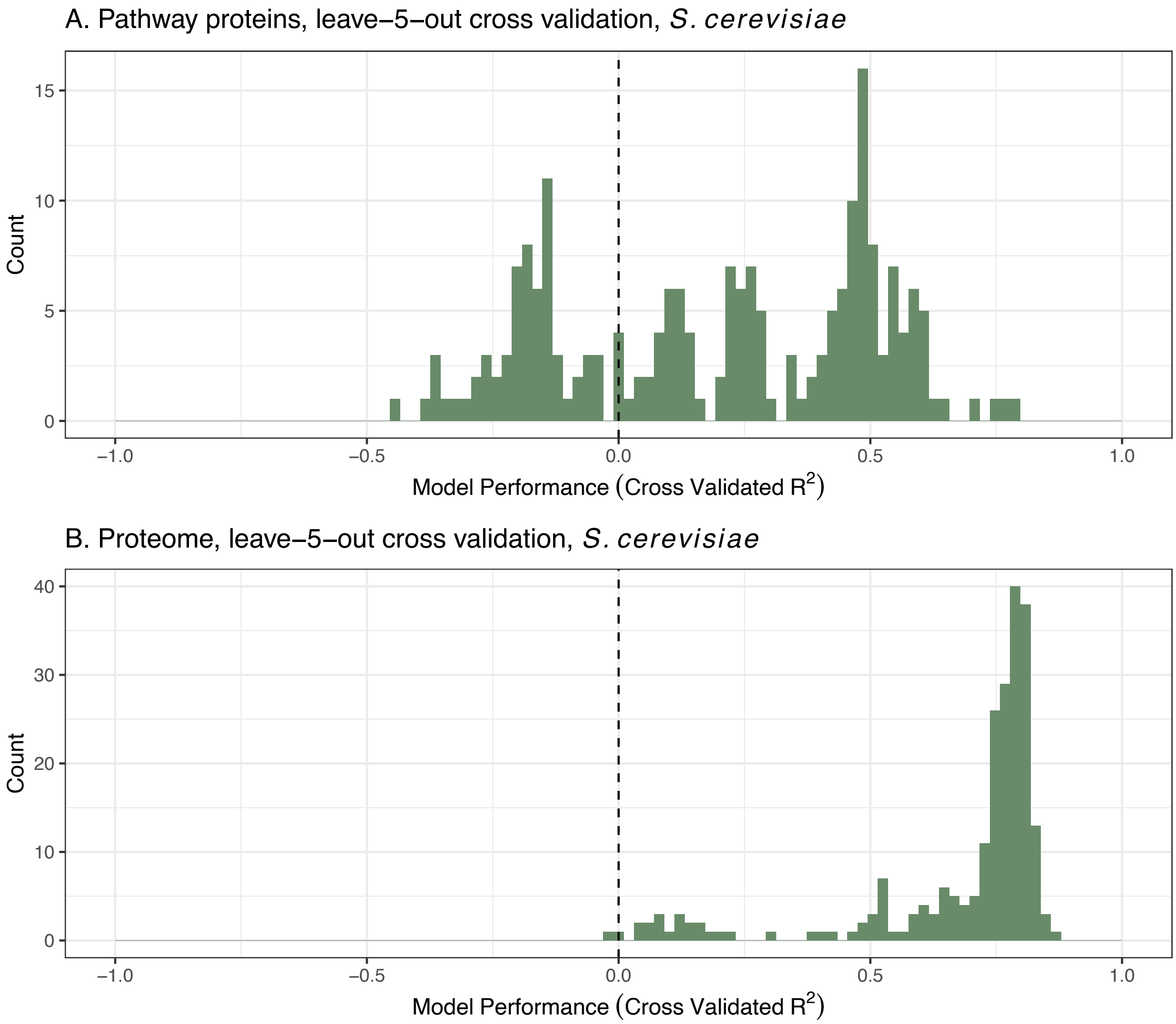


**Figure S8.** Distributions of cross-validated $R^2$ using leave-5-out cross-validation for the pathway level (A) and proteome level predictions (B). The specific model used was a ridge regression. There are 205 and 230 predictive model summaries in total for the top and bottom panels, respectively.

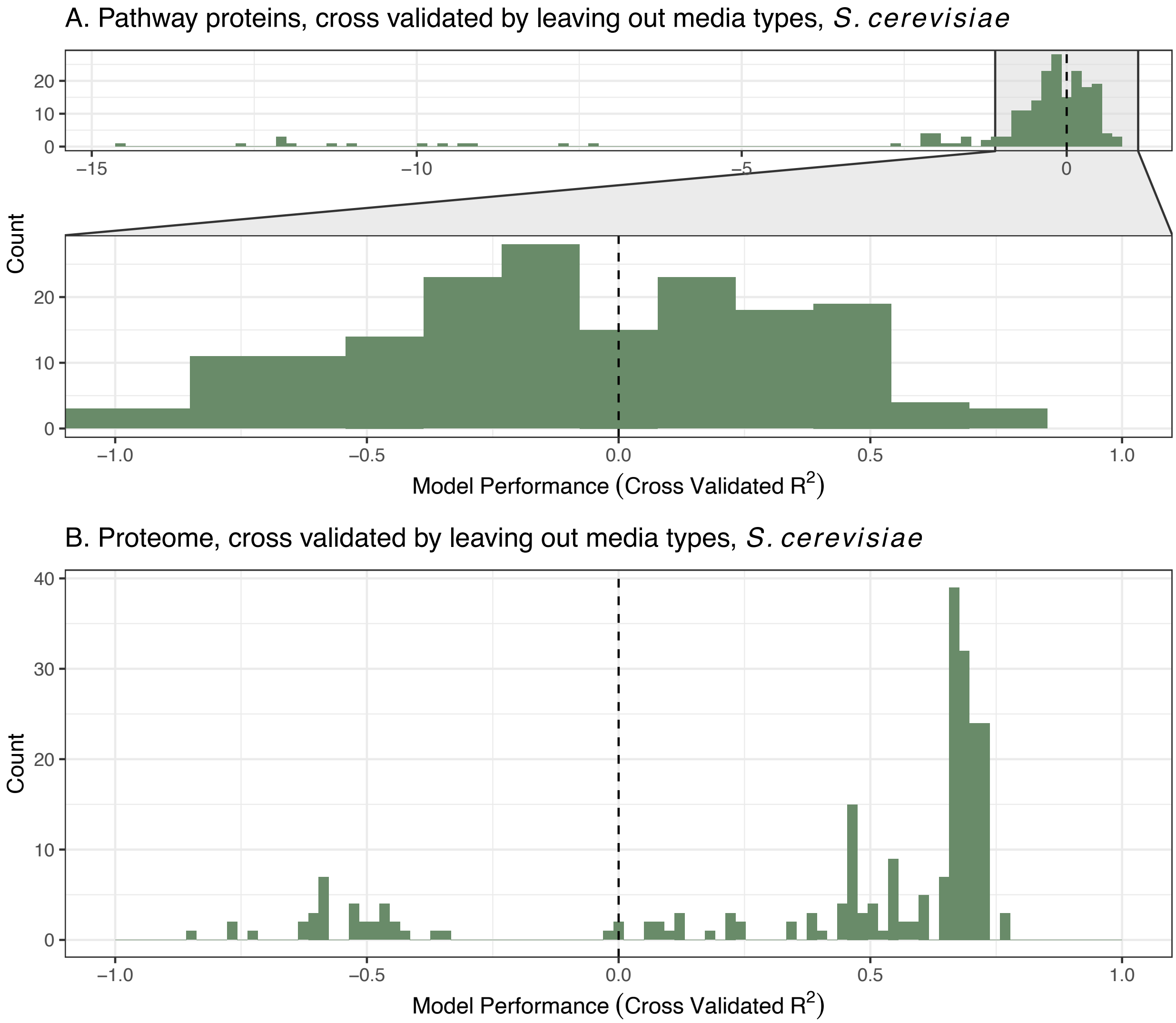


**Figure S9.** Distributions of cross-validated $R^2$ using block cross-validation across limiting substrate types (n = 5 different limiting substrates) for the pathway level (A) and proteome level predictions (B). The specific model used was a ridge regression. There are 205 and 230 predictive model summaries in total for the top and bottom panels, respectively.

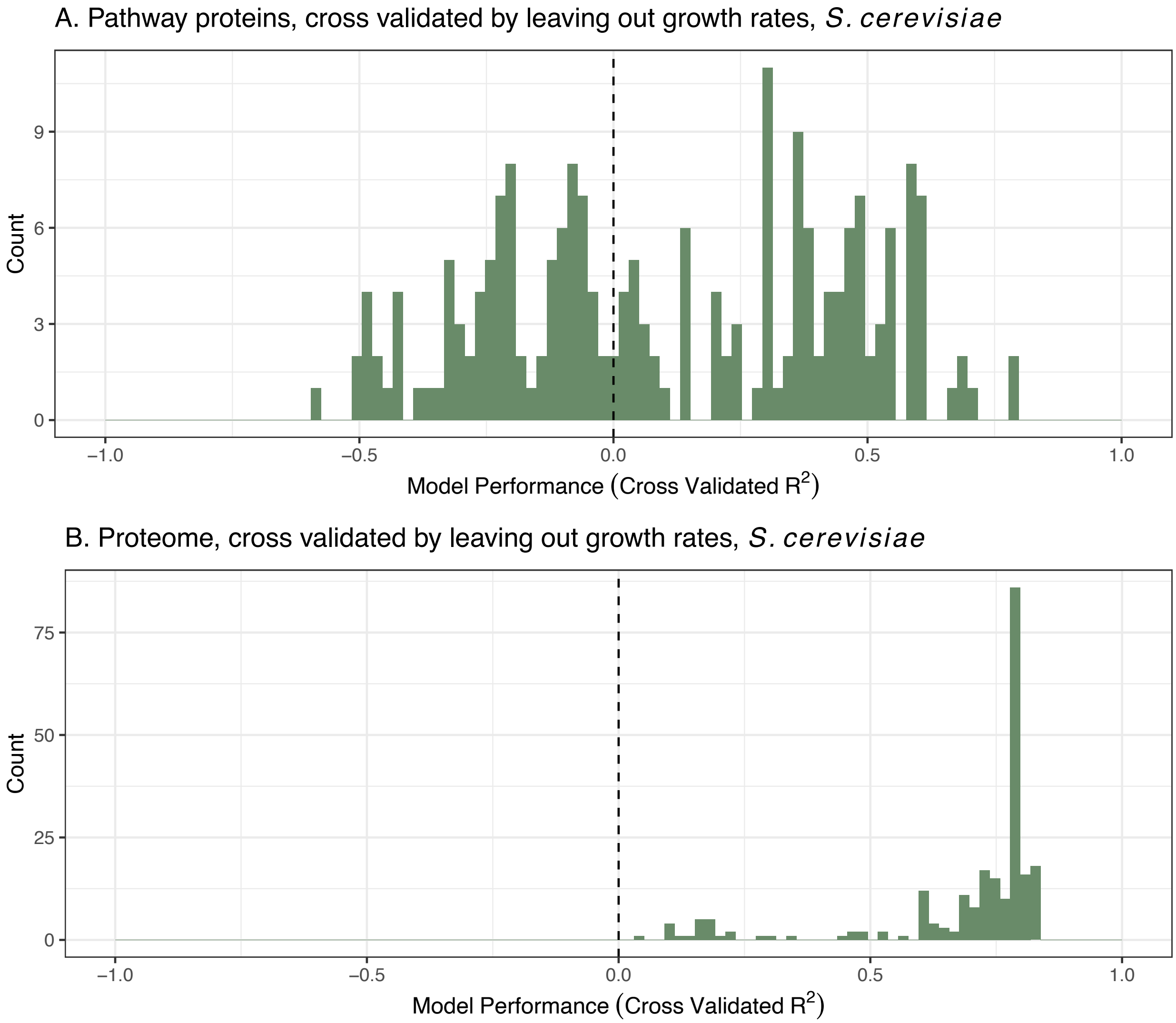


**Figure S10.** Distributions of cross-validated $R^2$ using block cross-validation across growth rates (dilution rates in the chemostat) for both the pathway level (A) and proteome level predictions (B), using ridge regressions. There are 208 and 233 predictive model summaries in total for the top and bottom panels, respectively.

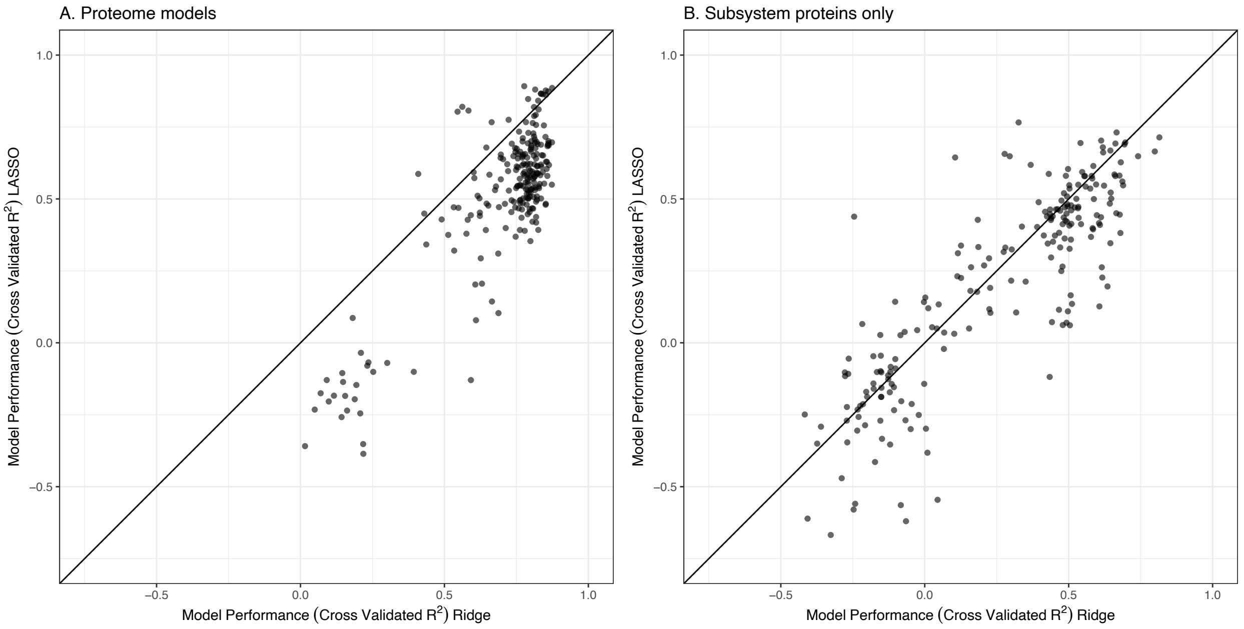


**Figure S11.** Comparison between leave-two-out cross-validated ridge regression $R^2$ with leave-two-out cross-validated LASSO regression $R^2$ for the (A) proteome-level models and the (B) subsystem-level models. There are 233 and 208 predictive model comparisons for panels A and B, respectively.

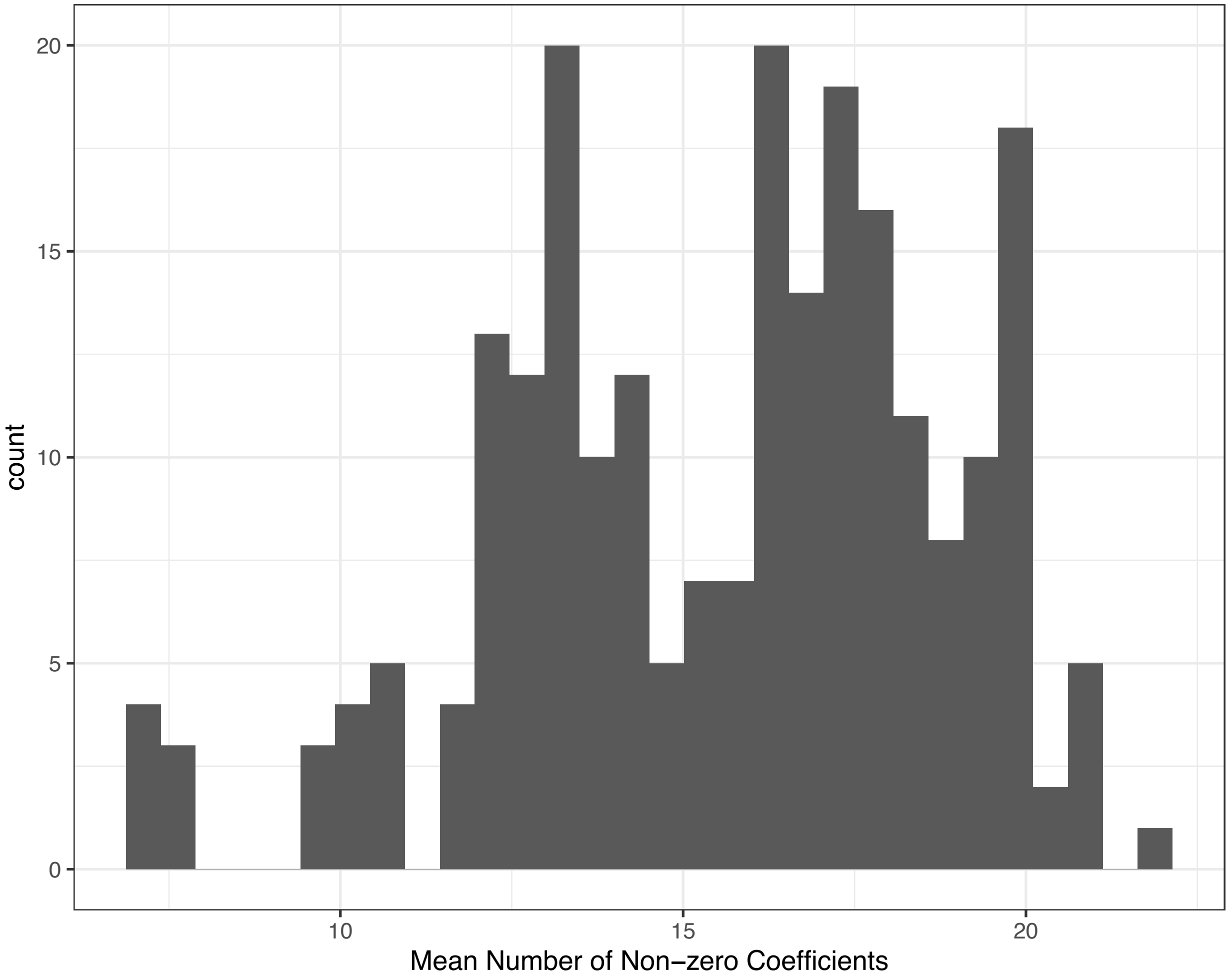


**Figure S12.** LASSO regression models result in a number of non-zero coefficients for predictive proteins. For each rate, we used cross-validated LASSO regression models to examine this number, and then calculated the mean number (across cross-validation iterations). This histogram (including 233 model summary statistics) shows the distribution of these mean values.

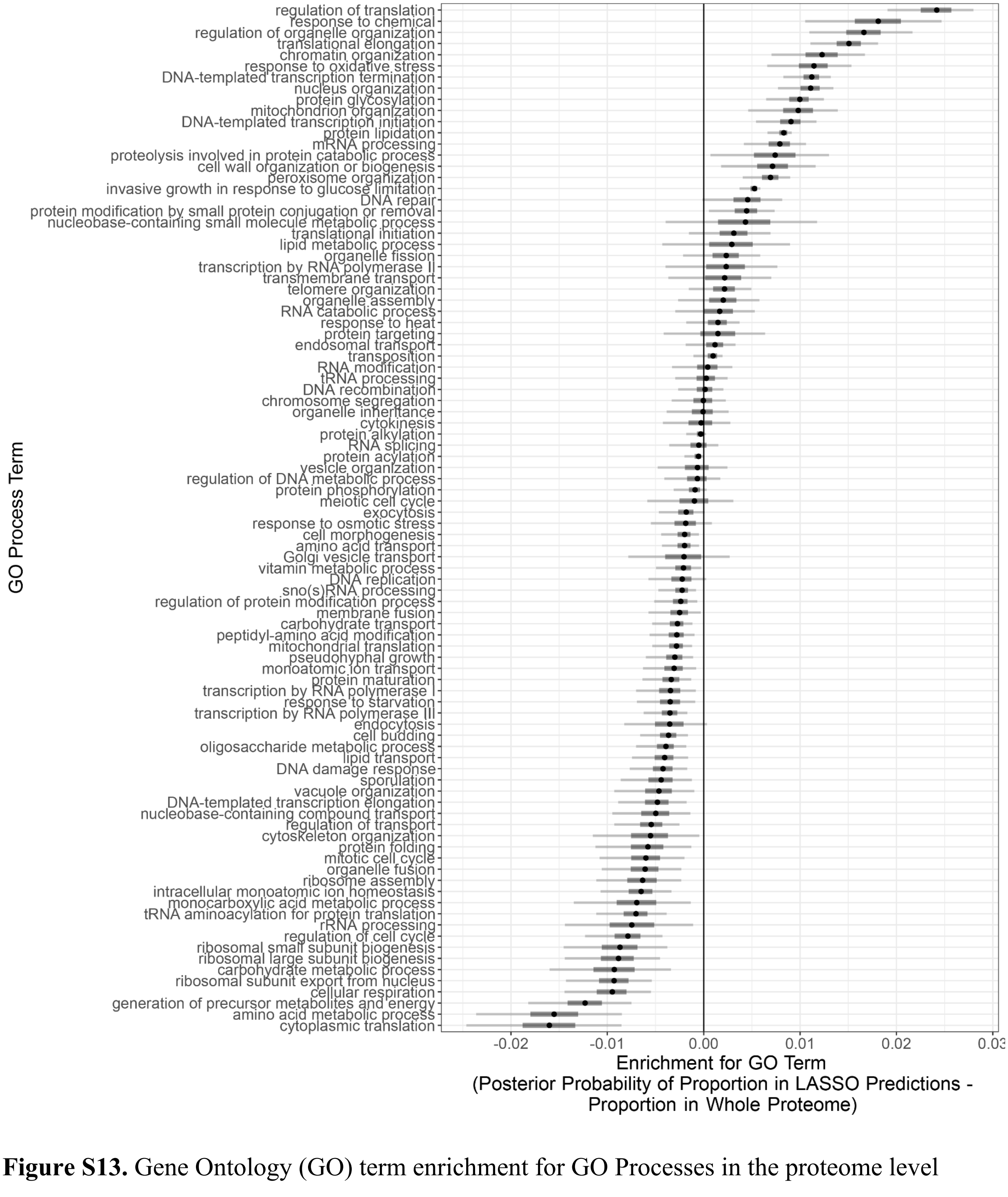


**Figure S13.** Gene Ontology (GO) term enrichment for GO Processes in the proteome level LASSO predictive models. Values above zero represent that proteins were more frequently included in predictive models compared with the whole set of observed proteins. Values below zero represent proteins that were less frequently included in LASSO models. Bars represent the difference between posterior probability distributions for the estimated proportion in LASSO predictions minus the estimated proportion in the whole proteome. Points are the distribution median, thicker bars are the 25th and 75th quantiles, and thin bars are the 2.5th and 97.5th quantiles. Note that the uncertainty is also a function of the number of cross-validation iterations for the LASSO model, so we only use it here as a heuristic.

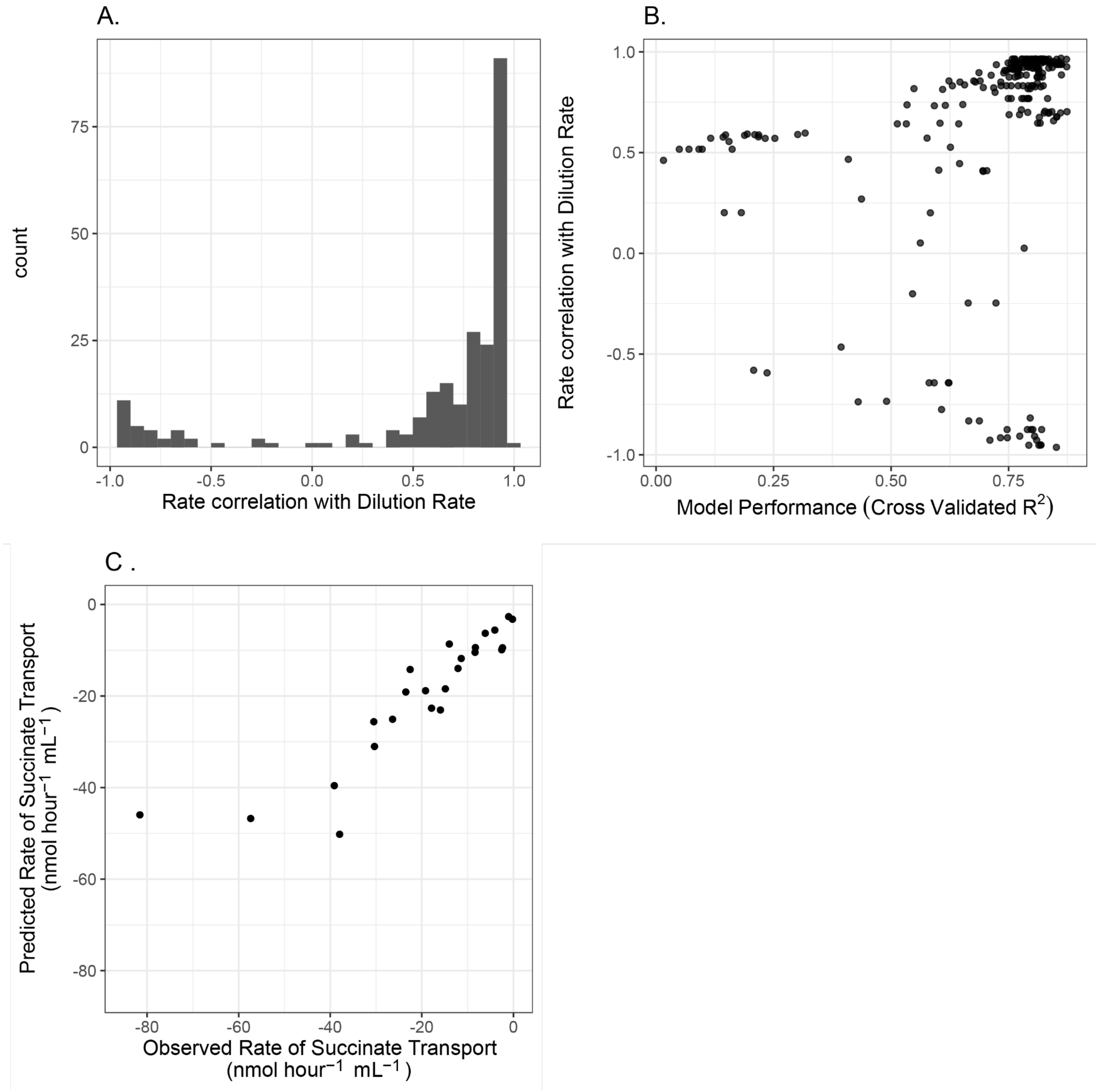


**Figure S14.** (A) The distribution of reaction rate to dilution rate Pearson correlation coefficients (233 values). (B) Relationship between model performance (assessed with cross-validated $R^2$) and the rate correlation with growth rate (233 points). (C) Succinate transport predictions using proteome-wide data versus observed succinate transport (shown is leave-two-out cross-validation predictions; 25 predictions).
